# Kinesin-1 mediates proper ER folding of the Ca_V_1.2 channel and maintains mouse glucose homeostasis

**DOI:** 10.1101/2024.06.24.600327

**Authors:** Yosuke Tanaka, Atena Farkhondeh, Wenxing Yang, Hitoshi Ueno, Mitsuhiko Noda, Nobutaka Hirokawa

## Abstract

Glucose-stimulated insulin secretion (GSIS) from pancreatic beta cells is a principal mechanism for systemic glucose homeostasis, of which regulatory mechanisms are still unclear. Here we show that kinesin molecular motor KIF5B is essential for GSIS through maintaining the voltage-gated calcium channel Ca_V_1.2 levels, by facilitating an Hsp70-to-Hsp90 chaperone exchange to pass through the quality control in the endoplasmic reticulum (ER). Phenotypic analyses of KIF5B conditional knockout (cKO) mouse beta cells revealed significant abolishment of glucose-stimulated calcium transients, which altered the behaviors of insulin granules via abnormally stabilized cortical F-actin. KIF5B and Hsp90 colocalize to microdroplets on ER sheets, where Ca_V_1.2 but not K_ir_6.2 is accumulated. In the absence of KIF5B, Ca_V_1.2 fails to be transferred from Hsp70 to Hsp90 and is likely degraded via the proteasomal pathway. KIF5B and Hsc70 overexpression increased Ca_V_1.2 expression via enhancing its chaperone binding. Thus, ER sheets may serve as the place of KIF5B- and Hsp90-dependent chaperone exchange, which predominantly facilitates Ca_V_1.2 production in beta cells and properly enterprises GSIS against diabetes.

## Introduction

Diabetes is a major metabolic syndrome, which is predicted to have a > 50% worldwide prevalence by 2045 (Perreault Skyler & Rosenstock, 2021). Glucose-stimulated insulin secretion (GSIS) is one of the central processes of systemic glucose homeostasis (Matschinsky, 1996). This process is regarded as a major target for therapeutic approaches to diabetes, so that detailed elucidation of stimulation-secretion-coupling is very important (Islam, 2020). The insulin secretion process largely consists of basal and glucose-stimulated secretion, the latter of which is classified into the first phase during 0–10 min and the second phase later than 10 min after the glucose stimulation (Wang & Thurmond, 2009). Basal insulin secretion is normally suppressed by hyperpolarization of the membrane potential, mainly due to K_ATP_ channel activity, which is augmented by the PIP_2_ synthesizing enzyme, PIP5Kα (de la Cruz *et al*., 2016; Liang *et al*., 2014). Na/K ATPase and voltage- and calcium-sensitive big K channel (BK_Ca_) contribute to this process as well, and circumvent beta cell hyperplasia (Arystarkhova *et al*., 2013; Dufer *et al*., 2009; Fridlyand *et al*., 2013). For the regulated secretion, glucose stimuli depolarize the membrane due to the K_ATP_ channel closure (Rorsman & Ashcroft, 2018), and induce Ca^2+^ influx mainly via the L-type voltage-gated calcium channel (VGCC) subunit Ca_V_1.2 and the R-type subunit Ca_V_2.3 (Rutter, 2001; Wilson *et al*., 2001). This Ca^2+^ influx is accompanied by glucose-stimulated sequential activation of Src-family kinases (SFKs) and the Rho-family GTPases Cdc42 and Rac1, which remodel subplasmalemmal F-actin bundles and stabilize insulin granule exocytosis events especially in the second phase (Arous & Halban, 2015; Daniel *et al*., 2002; Kalwat & Thurmond, 2013; Wang & Thurmond, 2010; Yoder *et al*., 2014). These Rho-family GTPase pathway may synergize with Ca^2+^-induced B-Raf/Raf1 pathways to activate PAK1/ERK kinases that directly drive the actin remodeling (Kalwat & Thurmond, 2013). Detailed molecular mechanism and the degree of involvement of Ca^2+^ in each glucose-stimulated signaling component is still largely elusive (Komatsu *et al*., 2013; Shigeto *et al*., 2006).

KIF5B is the founding member of kinesin superfamily proteins (KIFs)(Hirokawa *et al*., 2009). This ubiquitously expressed heavy chain of conventional kinesin-1 molecular motor was first cloned from mouse pancreatic beta cell cDNA library as a “beta cell kinesin” (Meng *et al*., 1997). Mice with complete knockout of this molecule are embryonic lethal (Tanaka *et al*., 1998). KIF5B plays an essential role in the second phase of GSIS (Cui *et al*., 2011; Donelan *et al*., 2002; Meng *et al*., 1997; Varadi *et al*., 2002; Varadi *et al*., 2003), but the precise molecular mechanism is still unclear. It transports various kinds of membrane organelles including mitochondria and lysosomes (Tanaka *et al*., 1998) and AMPA-type glutamate receptor-containing vesicles (Setou *et al*., 2002). This molecule also pulls out a tubular structure from the ER cisternae in vitro (Wozniak & Allan, 2006), and distributes the ER exit sites throughout the cytoplasm (Gupta *et al*., 2008). However, the relevance of these KIF5B-mediated processes in the functional regulation of protein folding in the ER are still largely unknown.

The ER is the location for the folding of de novo synthesized proteins (Braakman & Hebert, 2013). Being translated in the rough ER, membrane proteins undergo the folding process assisted by the ER-resident luminal chaperones calnexin-1 and derlin-1; and by the heat shock proteins (HSPs) that largely behave as cytoplasmic chaperones (Freeman & Morimoto, 1996; Yahara *et al*., 1998). In particular, some ER client proteins need a process termed “chaperone exchange,” in which the client-binding Hsp70 chaperone is replaced by the Hsp90 chaperone with the help of the STIP1 (Hop) protein (Donnelly *et al*., 2013; Moran Luengo *et al*., 2018), as well as the ER resident chaperones calnexin-1 and derlin-1 (Ramos *et al*., 2007; Volpi *et al*., 2016). Because this process plays an essential role in ER quality control (ERQC), the proteins that cannot undergo chaperone exchange are degraded by ER-dependent protein degradation (ERAD) involving the proteasomal pathway (Hoseki *et al*., 2010). Thus, the chaperone exchange is a critical checkpoint of protein folding for a subset of ER clients. However, its relationship to specific ER morphological compartment and kinesin molecular motor is still unclear.

In this study, we have investigated the molecular roles of KIF5B in beta cells in detail using the beta-cell-specific KIF5B conditional knockout (cKO) mice and KIF5B-knockdown MIN6 insulinoma cells. We propose that KIF5B is a crucial factor for Hsp70-to-Hsp90 chaperone exchange for Ca_V_1.2 acting on ER sheets and thus essentially maintains Ca_V_1.2 expression level that is indispensable for Ca^2+^-mediated GSIS for glucose homeostasis. This process involves Ca^2+^-mediated SNARE complex formation as well as Ca^2+^-dependent cortical F-actin remodeling, which allows insulin granules for short-range directional movements and full-fusion exocytosis. We also showed that K_ATP_ subunits can be folded irrespective of KIF5B and Hsp90, suggesting a selective role of the KIF5B–Hsp machinery for a subset of membrane protein synthesis. As we present that dual overexpression of KIF5B and Hsc70 in MIN6 cells results in upregulation of Ca_V_1.2 protein levels, KIF5B–Hsp machinery may serve as an important regulatory system that enhances the GSIS performance of beta cells, upon changing the systemic nutrition balance of the body.

## Results

### Beta-cell-specific KIF5B deletion in mice leads to glucose intolerance

We generated pancreatic beta-cell-specific conditional knockout mice for the *Kif5b* gene (*Kif5b^flox/flox^ Rip2-Cre*; cKO) using homologous recombination in mouse ES cells (Fig. EV1). The significant deficiency of KIF5B in insulin-expressing islet beta cells was verified by immunohistochemistry (Fig. 1A). The cKO islets revealed significant hyperplasia (Fig. 1B and C), which may partly compensate the defect in GSIS.

**Figure 1.**
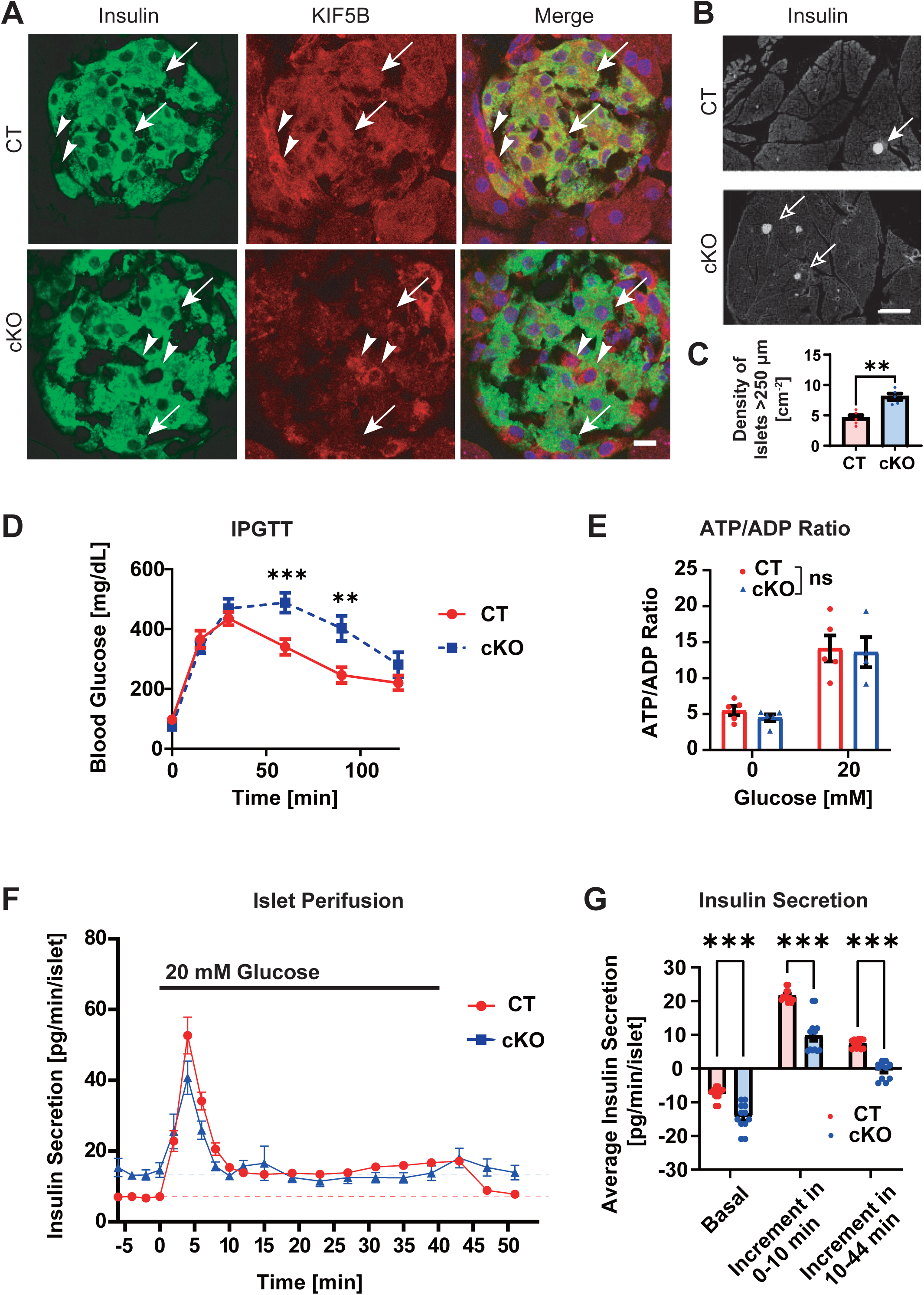
KIF5B is essential for proper GSIS and blood glucose homeostasis. (A–C) Immunohistochemistry of *Kif5b^flox/flox^ Rip2-Cre^●/●^* (CT) and *Kif5b^flox/flox^ Rip2-Cre^tg/●^* (cKO) mouse pancreas against insulin (green in A and gray in B) and/or KIF5B (red in A; A and B), accompanied by the islet morphometry (C). Bars, 10 μm in A and 1 mm in B. \*\**p* < 0.01, Welch’s *t* test, n = 5. Data are represented by the mean ± SEM throughout the panels. Arrows in A, beta cells. Arrowheads in A, non-beta cells. Arrows in B, islets. Corresponding to Fig. EV1. (D) Intraperitoneal glucose tolerance test (IPGTT) of 4-month-old CT and cKO mice. \*\*\**p* < 0.001; \*\**p* < 0.01; Welch’s *t* test, n = 4–5 at each time point. (E) ATP/ADP ratio measurements in the islets of the indicated genotypes stimulated for 3 min. NS, *p* > 0.05, n = 13–16, two-way ANOVA. (F and G) Perifusion assay of CT and cKO mouse pancreatic islets stimulated with 20 mM glucose for 45 min (F), quantified for the respective amounts of basal secretion (plotted in an inverted manner) and the first- (0–10 min) and second-phase (10–44 min) of GSIS increments. \*\*\**p* < 0.001; Welch’s *t* test, n = 12.

In the intraperitoneal glucose tolerance test (IPGTT), the cKO mice exhibited significant glucose intolerance compared with the floxed control (*Kif5b^flox/flox^*; CT; Fig. 1D). Because the isolated islets did not show significant alteration in the degree of elevating the ATP/ADP ratio after glucose stimulation (Fig. 1E), the metabolic state of glucose was supposed to be largely intact in cKO islets.

To investigate temporal transition in GSIS, we performed perifusion experiments of isolated islets (Fig. 1F and G). Interestingly, the basal level of insulin secretion in cKO islets was significantly elevated twice that in CT islets. In contrast, the peak size of the first-phase GSIS significantly decreased half of the control. The second-phase GSIS was further significantly abolished. These data were largely consistent with the results of previous genetic studies (Cui *et al*., 2011; Donelan *et al*., 2002; Meng *et al*., 1997; Varadi *et al*., 2002; Varadi *et al*., 2003), as well as the results of our preliminary study using aged *Kif5b^+/^*^-^ mice and their islets, so the consequence of genetic artifacts could be largely neglected.

### KIF5B is essential for full-fusion insulin granule exocytosis

To analyse the relevance of KIF5B in quantum events of exocytosis, we transduced primary culture of beta cells with synapto.pHluorin (Miesenbock *et al*., 1998; Tsuboi & Rutter, 2003), and observed them by TIRF microscopy between 30 and 60 min after glucose stimulation in the period of second phase GSIS (Fig. 2A and Movie EV1). The membrane fusion events were classified into kiss-and-run fusion events (< 1.2 s) and full-fusion events (≥ 1.2 s) according to previous insights (Ohara-Imaizumi *et al*., 2007; Takahashi *et al*., 2004; Takahashi *et al*., 2002) that is consistent with our observation with histogram analysis (Fig. 2B–D). Interestingly, cKO beta cells tended to exhibit more kiss-and-run fusion events but significantly fewer full-fusion events (Fig. 2E).

**Figure 2.**
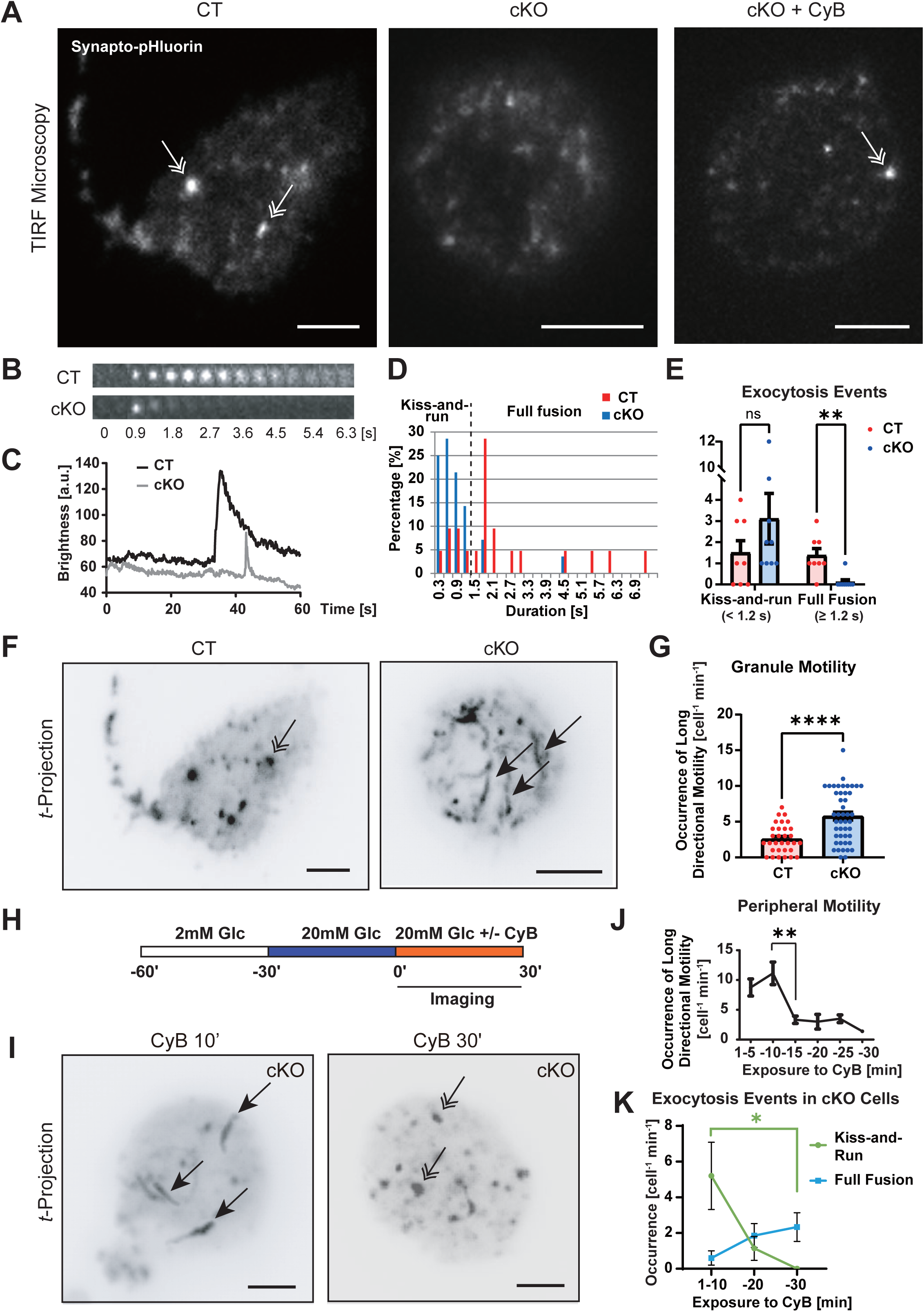
KIF5B stabilizes insulin exocytosis through actin remodeling. (A) Typical images of the surface of synapto.pHluorin-transduced primary beta cells of the indicated conditions in TIRF microscopy. Double arrows, full fusion. CyB, 10 µg/mL cytochalasin B treatment. Scale bars, 5 μm. Corresponding to Movie EV1. (B) Typical time-lapse images of insulin exocytosis on the cell surface of the indicated genotypes by time-lapse total internal reflection fluorescence (TIRF) microscopy, recorded at 30–60 min after glucose stimulation. (C) Typical traces of fluorescence intensities at the exocytosis points on the plasma membrane of the indicated genotypes. (D) Histogram of the duration of each exocytosis event at 30–60 min of glucose stimulation. (E) Quantification of the occurrence of full-fusion events (defined to be longer than 1.2 s in Panel D) and kiss-and-run events (shorter than 1.2 s) in each genotype. Note that KIF5B deficiency significantly reduced the exocytosis duration. ns, *p* > 0.05; \*\**p* < 0.01, Welch’s *t* test, n = 8–9. (F and G) Temporal projection of the time-lapse images of CT and cKO primary beta cells (F) and its quantification for the long directional motility (G). Scale bars, 5 μm. \*\*\*\**p* < 0.0001, Welch’s *t* test, n = 31–46. Double arrow, full fusion. Single arrows, cortical long-range directional motilities. (H–K) Time course of pharmacological rescue of insulin granule dynamics of cKO cells by CyB treatment, represented by an experimental design (H); temporal projection of time-lapse images (I); time course of the occurrence of peripheral directional motility after CyB treatment (J), and time course of the occurrence of indicated types of exocytosis after CyB treatment (K). Scale bars, 5 μm. \**p* < 0.05, \*\**p* < 0.01, compared with the data at 1–10 min, Welch’s *t* test, n = 4–8 (J and K).

By examining the *t-*projection views, we found that insulin granules in cKO beta cells significantly exhibited long-range movements passing through the fusion site, which reduced the duration of fusion events (Fig. 2F and G).

Because F-actin remodeling is known to stabilize full fusion exocytosis (Eitzen, 2003), we challenged cKO cells with a low dose (10 μg/mL) of the F-actin depolymerizing drug, cytochalasin B (CyB; Fig. 2H–K). Consequently, cKO cells treated by CyB for more than 10 min showed significantly and progressively fewer peripheral motility and kiss-and-run fusion events, and a greater occurrence of full-fusion events. Accordingly, KIF5B’s role in suppressing long-range directional transport of insulin granules for full fusion may be related with cortical actin remodeling.

### KIF5B allows directional short-range movements of insulin granules

To further investigate the dynamics of insulin granules, we labeled the granules of primary beta cells with phogrin-Dronpa-Green1 and subjected them to TIRF/PALM microscopy (Fig. 3A). The cortical density of insulin granules was largely unaltered by KIF5B deficiency (Fig. 3B).

**Figure 3.**
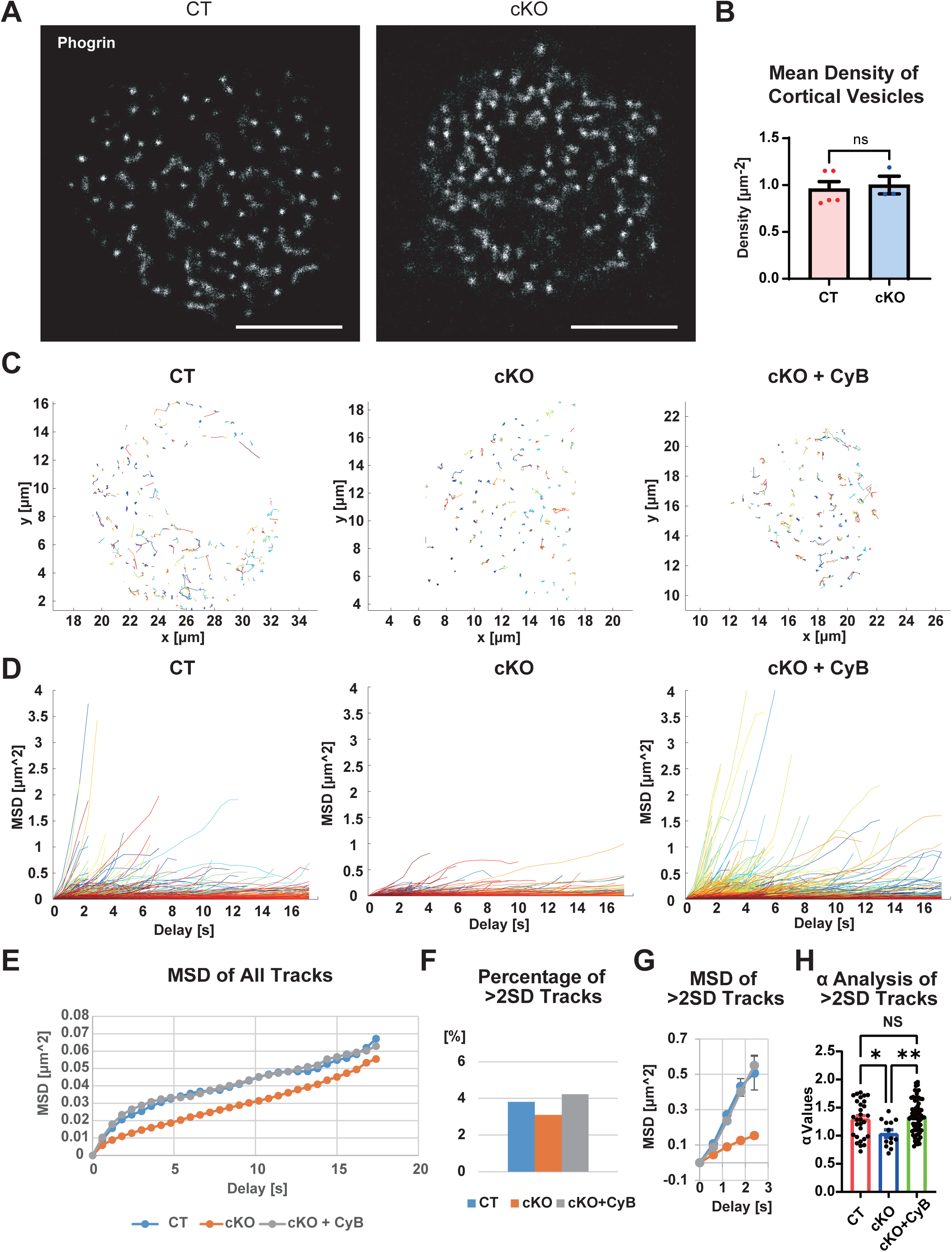
KIF5B facilitates short directional motility of insulin granules. (A and B) Images of cortical insulin granules of control (CT) and cKO primary beta cells, transfected with a *phogrin-Dronpa-Green1* expression vector, starved, and stimulated by 20 mM glucose for 30 min (A), accompanied by quantification of cortical insulin granules (B). Bar, 2 µm. ns, *p* > 0.05, n = 3–5, Welch’s *t* test. (C–H) Quantification of insulin granule motility tagged by phogrin-EGFP with a 5LIVE-Duo confocal microscope, in primary beta cells of the indicated genotypes and treatments for 100 s, at 30 min after glucose stimulation, represented by particle tracks (C), MSD trajectories (D), MSD curves of all tracks (E), percentage of >2 SD tracks (F), MSD curves of >2 SD tracks (G), and α analysis of >2 SD tracks for directional movements (H); corresponding to Movie EV2. Scale bar, 2 µm. Color coding in C and D, the time sequence. Note that the cKO beta cell granules were significantly less motile than the CT beta cell granules, which was significantly reversed by the CyB treatments.

Then, we labeled them with phogrin-EGFP and performed high-speed confocal imaging using an LSM 5LIVE-Duo microscope followed by particle tracking (Movie EV2). In this way, we detected loss of short-range directional movements of insulin granules in cKO beta cells (Fig. 3C), which again tended to be restored by CyB treatment.

These data were further subjected to mean square displacement (MSD) analyses (Fig. 3D–H), according to previously described methods (Tarantino *et al*., 2014). In the summary of individual trajectories, the slopes of CT and cKO + CyB ones were apparently steeper than cKO trajectories (Fig. 3D). The curves of MSD of all tracks indicated that the ones for CT and cKO + CyB were similar but the one for cKO was lower than those, mainly because of the existence of a convex upward component in 0–3 s of delay (Fig. 3E).

To further characterize this component, we depicted > 2 SD trajectories (Fig. 3F) and again summarized them, which revealed more significant changes in the slopes (Fig. 3G). Finally, we calculated the α values of each > 2 SD trajectory (Fig. 3H), where α = 1 represents freely diffusive movements and the higher values represent the existence of active transport. Very interestingly, the CT and cKO + CyB groups had similar average values of 1.3 but the cKO group had an average value of 1.0. Thus, KIF5B was suggested to enable active short-range actomyosin-based transport of insulin granules that may serve for the perpendicular access of granules to plasma membrane as previously observed in other systems (Ueno *et al*., 2011). Because CyB treatment could nicely compensate the short-range directional motilities of insulin granules, KIF5B’s role in enhancing short-range directional transport of insulin granules for full fusion was also considered to be related with cortical actin remodeling.

### KIF5B deficiency impairs F-actin remodeling during the second phase

Then we investigated whether glucose-stimulated F-actin remodeling is really disorganized in KIF5B-deficient cKO cells. In starved cells, the intensity of cortical F-actin was largely unaltered (Non-Stim; Fig. 4A). However, glucose stimulation for 60 min revealed a difference (Glucose; Fig. 4A). In CT cells, cortical F-actin was significantly remodeled to become thinner during this period, as previously reported (Nevins & Thurmond, 2003; Olofsson *et al*., 2009). Because F-actin tended to accumulate into foci on the cell surface, mutual sliding and depolymerization may be involved in this remodeling process. In contrast, in the cKO cells, cortical F-actin was not remodeled but was even thickened by glucose stimulation. The relative median intensity of cortical F-actin of CT cells became half in this period, but that of cKO cells was almost unchanged (Fig. 4B). Because forced [Ca^2+^]_i_ elevation by ionomycin could restore the F-actin remodeling in both genotypes, the role of KIF5B in F-actin remodeling was considered to be dependent on or in parallel to Ca^2+^ transients.

**Figure 4.**
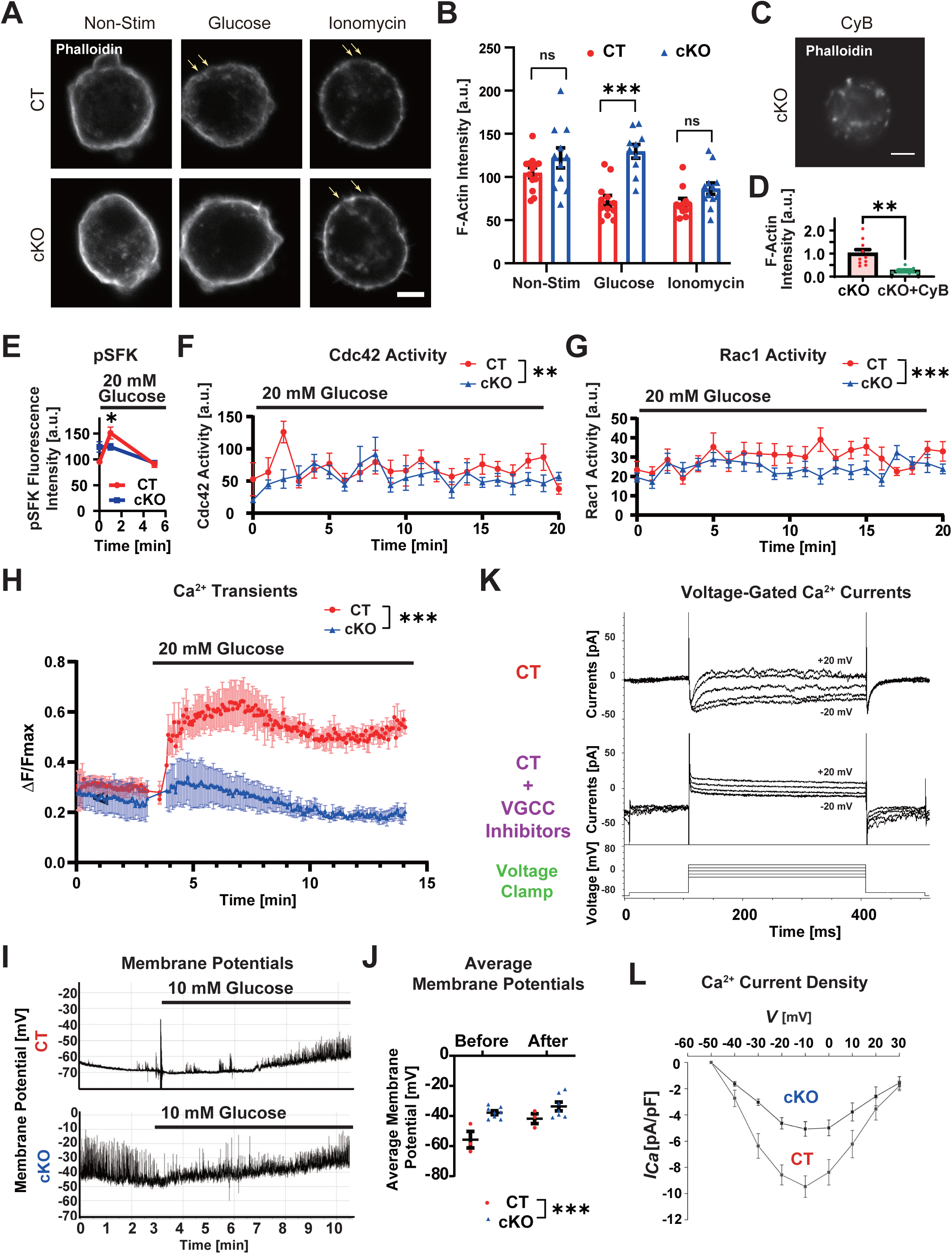
KIF5B is essential for Ca^2+^ transients of beta cells and actin remodeling. (A–D) Actin remodeling assay using fluorescent phalloidin-labeled CT and cKO primary beta cells with the indicated treatments (A and C), accompanied by quantification (B and D). Scale bar, 5 μm. ns, *p* > 0.05, \*\*\**p* < 0.001; Welch’s *t* test, n = 10–13. Quantified at the positions of largest diameter. Arrows, the sites of cortical F-actin remodeling. Corresponding to Fig. EV2A–D and Movie EV3. (E–G) Glucose-stimulated activation of SFK (E), Cdc42 (F), and Rac1 (G) in primary beta cells of the indicated genotypes after 20 mM glucose stimulation from time 0, measured by immunofluorescence microscopy (E) and the respective FRET biosensors (F and G). \**p* < 0.05, \*\**p* < 0.01, \*\*\**p* < 0.001; n = 28–42, Welch’s *t* test (E); and n = 22 (F and G), 2-way ANOVA (F and G). (H) 20 mM glucose-stimulated calcium transients of the primary beta cells labeled with Fluo4-AM. \*\*\**p* < 0.001, n = 3, two-way ANOVA on periods after the stimulation. (I and J) Membrane potentials of resting and 10 mM glucose-stimulated primary beta cells according to whole-cell patch-clamp recordings (I); and quantification of the mean membrane potentials before and after the glucose stimulation (J). \*\*\**p <* 0.001, two-way ANOVA, n = 3–8. (K) Traces of voltage-gated Ca^2+^ currents in patch-clamp recording of mouse primary beta cells of the indicated genotypes in 10 mM glucose, with or without the VGCC inhibitors cocktail, containing 20 μM nifedipine, 1 μM SNX482, and 0.3 mM ascorbate, at the range of -20 to +20 mV. Note that the inhibitor treatment significantly abolished the voltage-gated inward currents. (L) Ca^2+^ inward current density curves of the primary beta cells of the indicated genotypes measured by whole-cell patch clamp.

This phenotype was reproduced in a time-lapse manner using primary beta cells of *Lifeact-mCherry* transgene-carrying mice (Fig. EV2A; Movie EV3). We conducted *Kif5b* gene silencing as described later. Scrambled-control miRNA-transduced (SC) cells exhibited gradual remodeling of peripheral actin bundles by 5–25 min after glucose stimulation. Interestingly, large pseudopods were dynamically produced in KIF5B knockdown (KD) cells, but peripheral F-actin remained intact upon glucose stimulation. As a control, KIF5B-EYFP overexpression in cKO cells also significantly restored cortical F-actin remodeling (Fig. EV2B–D), suggesting that these phenotypes were truly from KIF5B deficiency. The application of CyB during glucose stimulation completely disrupted and fragmented the cortical F-actin of cKO cells (Fig. 4C and D).

To study upstream signaling pathway leading to F-actin remodeling (Veluthakal & Thurmond, 2021), we measured the time course of the level of phosphorylated and activated SFK (pSFK) using immunofluorescence, and those of the Cdc42 and Rac1 GTPase activities using FRET biosensors (Yoshizaki et al, 2003). In CT primary beta cells, a transient SFK activation at 1 min, Cdc42 activation at 1–3 min, and a sustained Rac1 activation later than 5 min were sequentially observed (Fig. 4E–G). However, in cKO primary beta cells, these activities were significantly abolished. These data collectively indicated an essential role of KIF5B in F-actin remodeling through its relevance in SFK–Rho-family GTPase signal transduction cascade.

### KIF5B supports electrophysiological and Ca^2+^ activities in beta cells

Interestingly, glucose-induced Ca^2+^ transients were significantly impaired in cKO beta cells (Fig. 4H). In CT primary beta cells, glucose application significantly elevated the [Ca^2+^]_i_ immediately after application to reach the maximum in 3 minutes. However, in cKO cells, the level of glucose-stimulated [Ca^2+^]_i_ elevation was less than one-fifth that in CT cells.

Then we conducted patch-clamp recording of voltage-gated Ca^2+^currents in primary beta cells. First, we conducted whole cell patch clamp of primary beta cells with glucose elevation from 0 to 10 mM in the external solution (Fig. 4I and J). The resting membrane potential of CT cells was properly maintained at -60 mV. However, those of cKO beta cells were significantly depolarized to -40 mV. Consequently, spontaneous action potentials tended to increase in the resting state in cKO beta cells. This membrane excitation tendency can partly explain the cause of beta cell and islet hyperplasia (Li *et al*., 2014).

Then, voltage-gated Ca^2+^ currents were recorded in 10 mM glucose with stepwise changes in membrane potentials (Fig. 4K). The observed voltage-gated inward currents were suppressed by a VGCC inhibitor cocktail. Under this condition, the Ca^2+^ current densities in cKO beta cells were significantly smaller than those of CT beta cells (Fig. 4L). Accordingly, cKO beta cells exhibited continuous membrane excitation, while the voltage-gated inward Ca^2+^ current densities in 10 mM glucose were significantly reduced, leading to the abolishment in the glucose-stimulated Ca^2+^ transients.

### KIF5B deficiency leads to downregulation of membrane proteins including Ca_V_1.2

To investigate the molecular mechanism between KIF5B and Ca^2+^ transients, we conducted gene silencing by expressing *Kif5b-*antisense knockdown (KD) or scrambled sequence control (SC) miRNAs using mammalian expression vectors in MIN6 insulinoma cells (Fig. 5A and B). As a result, the levels of Ca_V_1.2, Ca_V_2.3, PIP5Kα, BK_Ca_, syntaxin-1, Na/K ATPase, and Hsp70 proteins were significantly reduced. In contrast, the K_ATP_ channel subunits K_ir_6.2 and SUR, as well as α-tubulin and calnexin-1 were largely unaltered. Because the difference in transcription levels of Ca_V_1.2, Ca_V_2.3, and K_ir_6.2 were subtle (Fig. 5C), the reduction appeared to occur primarily at the post-transcriptional level.

**Figure 5.**
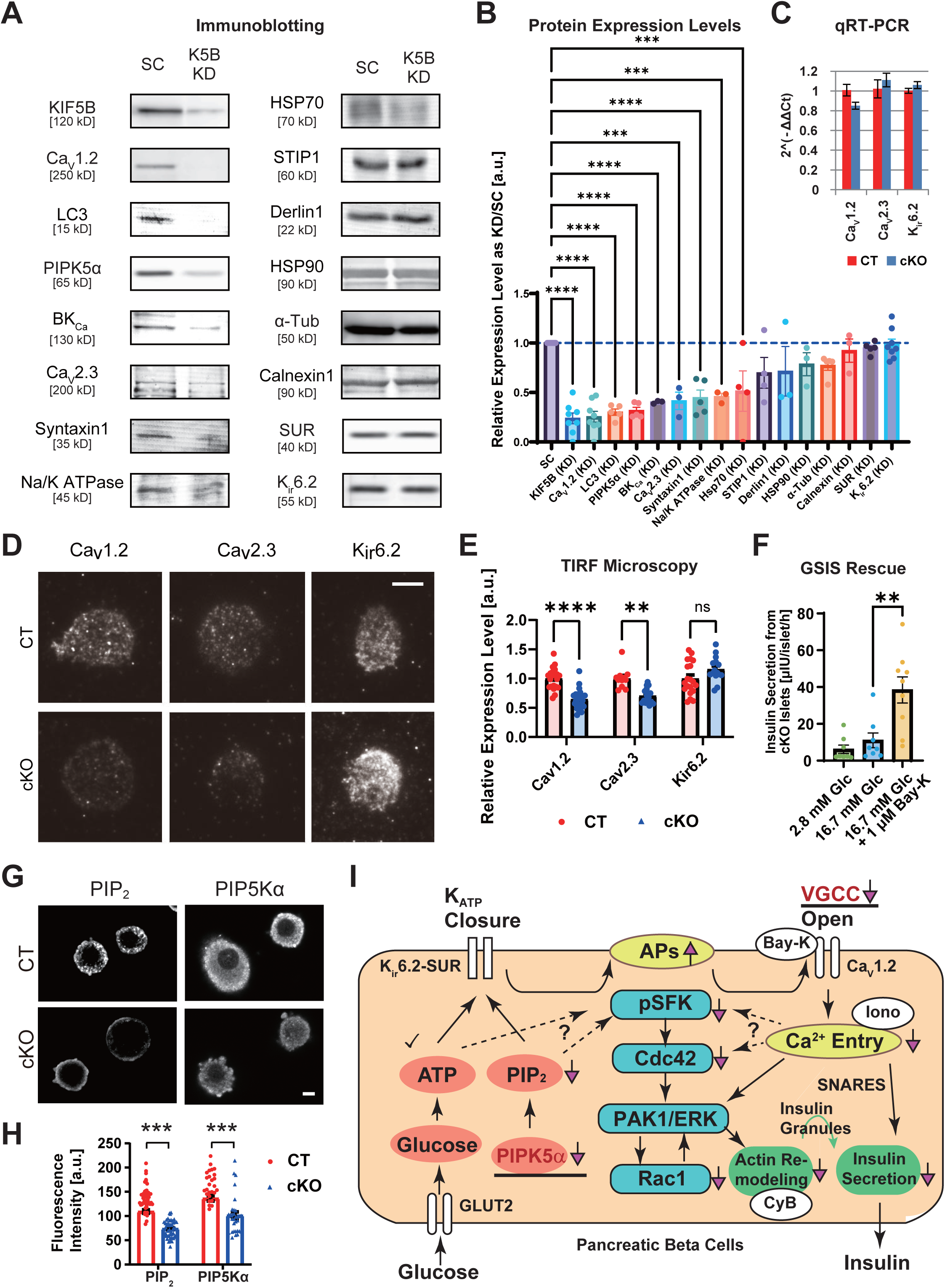
KIF5B is essential for Ca_V_1.2 protein expression in beta cells. (A and B) Immunoblotting of a *Kif5b* gene silencing system in MIN6 cells using scrambled control (SC) and KIF5B-knockdown (KD) miRNA, for the indicated proteins (A); and its normalized quantification against the SC-transduced cells (B). \*\*\**p* < 0.001; \*\*\*\**p* < 0.0001; Welch’s *t* test between KD and SC; n = 3–8. (C) Quantitative RT–PCR results of the KD system among the indicated genes. n = 6. (D and E) Immunofluorescence microscopy of primary beta cells of the indicated genotypes, against Ca_V_1.2, Ca_V_2.3, and K_ir_6.2, using TIRF microscopy (D), and its quantification (E). Bar, 5 μm. ns, *p* > 0.05; \*\**p* < 0.01; \*\*\*\**p* < 0.0001; n = 11–21, two-way ANOVA. (F) Rescue of impaired GSIS from cKO mouse islets by 1 μM Bay-K 8644. \*\**p <* 0.01, one-way ANOVA, n = 8–9. (G and H) Immunofluorescence microscopy of primary beta cells of the indicated genotypes, against PIP_2_ and PIP5Kα using a confocal laser-scanning microscope (CLSM, G), and quantification (H). Bars, 5 μm. \*\*\**p* < 0.001; n = 26–50, Welch’s *t* test. (I) Schematic representation of possible changes in the stimulation-secretion coupling of GSIS in KIF5B cKO beta cells. Checkmark, normal expression. Red arrows, changes according to KIF5B deficiency. Ca_V_1.2 and PIP5Kα protein downregulation (underlined) in KIF5B-deficient beta cells may primarily result in the abolishment of Ca^2+^ transients and downregulation of PIP_2_, respectively. White ovals, pharmacological reagents that directly stimulated the respective pathways: Bay-K, Bay-K 8644; Iono, ionomycin; CyB, cytochalasin B.

We then conducted immunofluorescence microscopy in primary beta cells. Ca_V_1.2 and Ca_V_2.3 were significantly downregulated from the surface of cKO cells in TIRF microscopy (Fig. 5D and E). However, K_ir_6.2 remained intact or even slightly increased. To investigate whether a Ca_V_1.2 activation can compensate the GSIS failure due to KIF5B deficiency, we incubated the cKO islets with high glucose and/or the L-type VGCC agonist, Bay-K 8644 (Satin *et al*., 1995). Although high glucose could scarcely stimulate the GSIS of cKO islets, application of 1 *μ*M Bay-K 8644 along with high glucose stimulation yielded 8-fold increase of GSIS (Fig. 5F). These data suggested that pharmacological activation of the reduced level of Ca_V_1.2 in cKO islets could still restore the GSIS. Thus, we considered that cKO mouse islets may preserve the capacity for GSIS if proper Ca^2+^ transients were formed, and that the reduced expression of Ca_V_1.2 in cKO islets could be a major cause of the GSIS failure. The negative effects of ‘cellular fatigue’ through constitutive depolarization (Khaldi *et al*., 2004; Pertusa *et al*., 2002; Roche *et al*., 1998) was thus unlikely especially in the case of this second phase GSIS.

In addition, PIP5Kα and its enzymatic product PIP_2_ were significantly downregulated in cKO beta cells (Fig. 5G and H). Because PIP_2_ is a key regulator of K_ATP_, VGCC, and focal adhesion kinase (FAK) activities (Lin *et al*., 2005; Suh *et al*., 2012; Zhou *et al*., 2015), this downregulation may partly explain the accessory phenotypes in KIF5B cKO mice. Those changes in the stimulation-secretion coupling of KIF5B deficient beta cells were schematically represented in Fig. 5I.

### Ca_V_1.2 undergoes Hsp70-to-Hsp90 chaperone exchange and is stabilized by KIF5B

Focusing on the behavior of the VGCCs, we compared protein stability using cycloheximide (CHX) treatment that inhibited the whole protein synthesis (Fig. 6A–C). Ca_V_1.2 and Ca_V_2.3 protein signals in cKO cells degraded significantly faster than those in CT cells, whereas the degradation rate of α-tubulin was unchanged. To determine whether this downregulation of Ca_V_1.2 was caused by aberrant post-Golgi anterograde trafficking, we conducted a brefeldin A (BFA) washout experiment (Fig. 6D and E, Movie EV4). The time needed for Cav1.2–EGFP to reach the plasma membrane after BFA washout in cKO cells was only slightly decreased. Thus, kinesin-1 is largely dispensable for post-Golgi trafficking of Ca_V_1.2, which cannot be the major cause of the Ca_V_1.2 downregulation in cKO cells.

**Figure 6.**
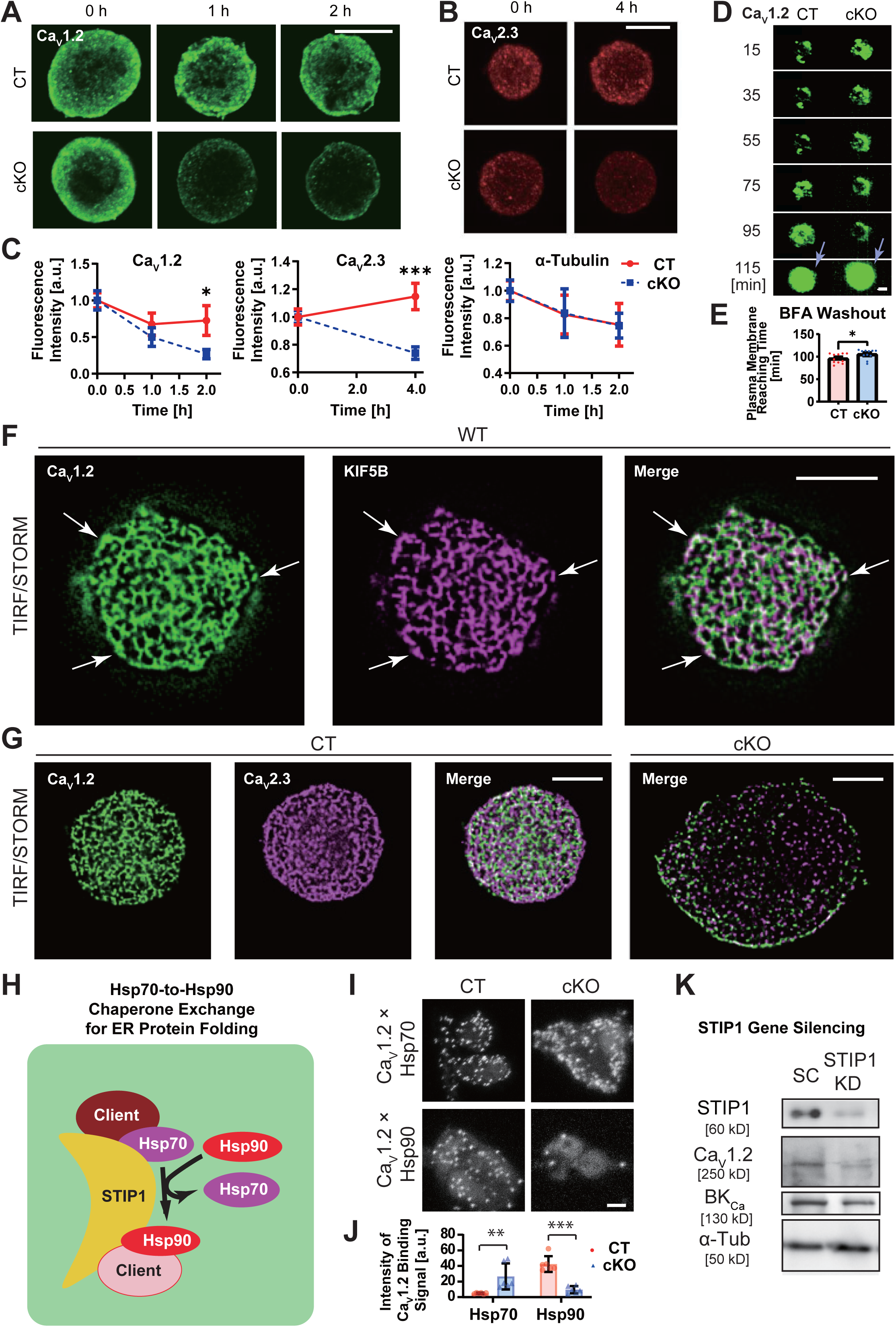
KIF5B facilitates chaperone exchange for Ca_V_1.2 protein expression. (A–C) Degradation assay in immunofluorescence against Ca_V_1.2 (A) and Ca_V_2.3 (B) of primary beta cells of the indicated genotypes after the CHX treatment of the indicated periods; accompanied by their quantification along with that of α-tubulin (C). Bars, 5 μm. \**p* < 0.05, \*\*\**p* < 0.001, Welch’s *t* test at the indicated time; n = 6 (Ca_V_1.2), 17–24 (Ca_V_2.3), 5–6 (α-tubulin). (D and E) Brefeldin-A (BFA) washout assay with an LSM 5LIVE-Duo microscope, assessing the speeds of post-Golgi trafficking of Ca_V_1.2-EGFP proteins expressed in primary beta cells of the indicated genotypes (D), accompanied by its quantification (E). Time after BFA washout is indicated. Bar, 5 μm. *\*p* < 0.05, Welch’s *t* test, n = 11. Arrows, the timing of plasma membrane fusion. Corresponding to Movie EV4. (F) TIRF/STORM microscopy of a wild-type primary mouse beta cell immunolabeled against Ca_V_1.2 and KIF5B. Scale bar, 5 μm. Arrows, colocalizing spots. (G) TIRF/STORM microscopy of primary mouse beta cells of the indicated genotypes immunolabeled against Ca_V_1.2 (green) and Ca_V_2.3 (magenta). Scale bars, 5 μm. (H) Schematic representation of STIP1-dependent Hsp70-to-Hsp90 chaperone exchange machinery. (I and J) *z*-projection of proximity ligation assay in CT and cKO primary beta cells showing the protein binding between Ca_V_1.2 and the indicated Hsp proteins (I); accompanied by quantification (J). \*\**p* < 0.01; \*\*\**p* < 0.001; Welch’s *t* test, n = 6. (K) Immunoblotting of scramble control (SC) and STIP1-knockdown (KD) MIN6 cells against the indicated epitopes. Note that STIP1 deficiency induced downregulation of Ca_V_1.2 and BK_Ca_ proteins. Reproduced twice.

We conducted TIRF/STORM microscopy of primary beta cells against KIF5B and Ca_V_1.2 (Fig. 6F). Very interestingly, these two proteins partially colocalized to ER-like networks beneath the plasma membrane of wild type cells. We also conducted TIRF/STORM microscopy for Ca_V_1.2 and Ca_V_2.3 in CT and cKO primary beta cells (Fig. 6G). These two VGCCs were also partially colocalized to ER-like network, and reduced by KIF5B deficiency. Thus, we sought to investigate the relevance of KIF5B in ER-mediated protein folding.

We focused on Hsp70-to-Hsp90 chaperone exchange in the middle way of ER protein folding (Fig. 6H), as kinesin-1 was previously identified to interact with Hsc70 in neuronal axons (Terada *et al*., 2010). We conducted a proximity ligation assay (PLA) to investigate the possible changes in chaperone binding capacities of Ca_V_1.2 protein in CT and cKO primary beta cells (Fig. 6I and J). Interestingly, KIF5B deficiency significantly affected the Ca_V_1.2–Hsp90 interaction, rather than Ca_V_1.2–Hsp70 interaction, suggesting its relevance in chaperone exchange of Ca_V_1.2.

STIP1 is an essential cochaperone for the chaperone exchange system (Bhattacharya & Picard, 2021; Bhattacharya *et al*., 2020; Harrison *et al*., 2024). We sought to investigate if STIP deficiency could affect the expression levels of the client protein candidates. Gene silencing of *Stip1* in MIN6 cells reproducibly resulted in decrease of Ca_V_1.2 and BK_Ca_ proteins as well as STIP1 protein itself (Fig. 6K), suggesting the involvement of chaperone exchange for the folding of at least these two proteins.

These data collectively suggested that KIF5B is required for chaperone exchange of a subset of ER clients including Ca_V_1.2.

### KIF5B expression facilitates chaperone binding capacities of Ca_V_1.2

To further characterize the cause of Ca_V_1.2 protein turnover in CHX-treated cKO beta cells, we applied the lysosomal inhibitor leupeptin or the proteasomal inhibitor MG-132 into the culture medium. In the consequence, MG-132, but not leupeptin, significantly restored the Ca_v_1.2 expression (Fig. 7A and B), suggesting the involvement of proteasomal system in its turnover, as reported in other systems (Altier *et al*., 2011).

**Figure 7.**
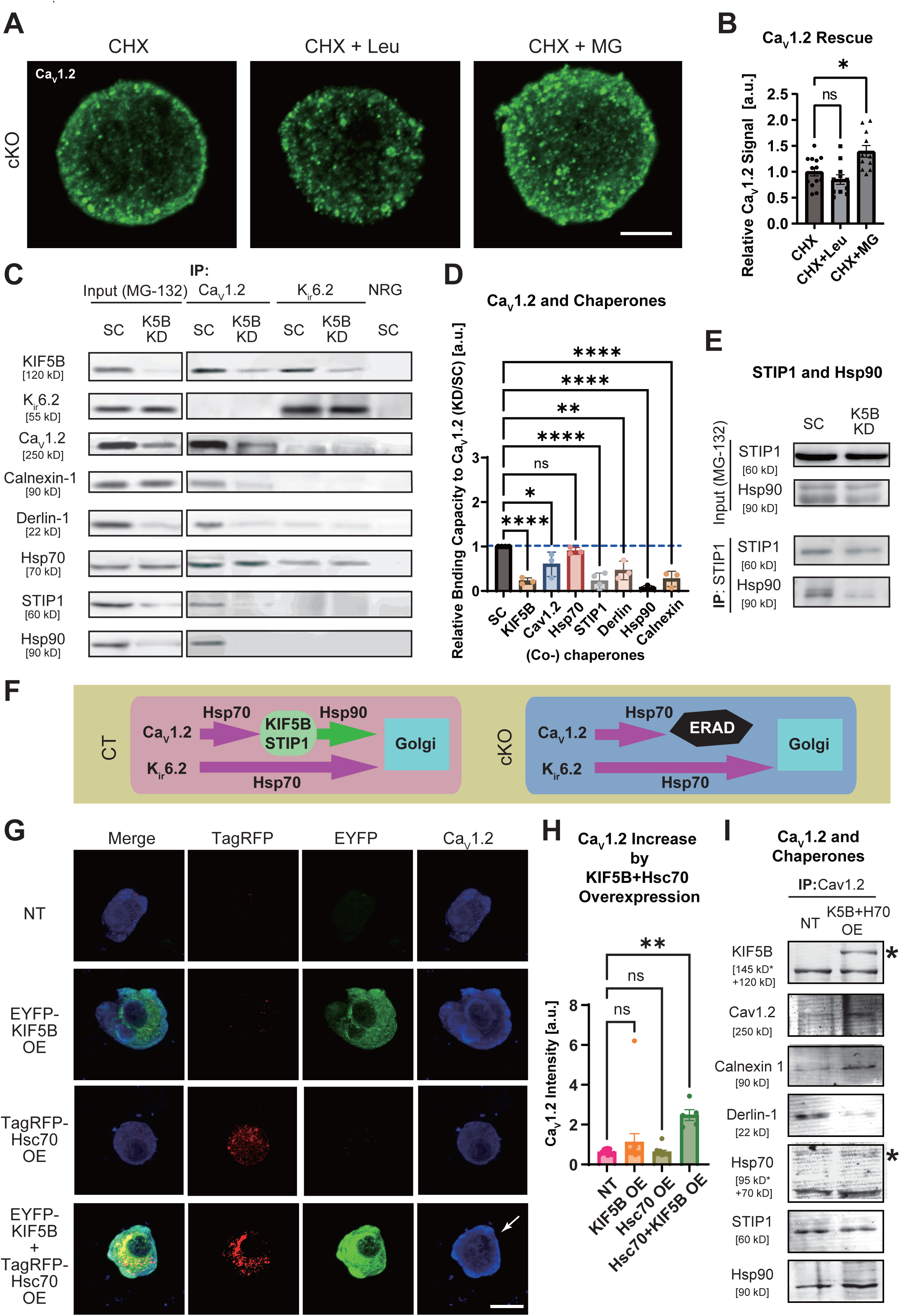
KIF5B–Hsp machinery positively regulates Ca_V_1.2 expression. (A and B) Rescue of Ca_V_1.2 degradation in cKO primary beta cells in the presence of CHX by leupeptin (Leu) or MG-132 (MG) for 4 h. ns, *p* > 0.05; \**p* < 0.05; one-way ANOVA, n = 12. (C and D) Vesicle IP of MG-132-treated MIN6 cell lysates transduced with scrambled control (SC) and KIF5B-knockdown (KD) miRNAs, precipitated using Ca_V_1.2 or K_ir_6.2 antibodies or normal rabbit IgG (NRG) and immunoblotted for the indicated proteins (C), accompanied by quantification of Ca_V_1.2-coprecipitated fractions (D). Note that the Ca_V_1.2-binding capacities of derlin-1, calnexin-1, and Hsp90 chaperones and that of the adaptor protein STIP1 in KD cell lysates were significantly lower than those in SC cell lysates. (E) Vesicle IP of the MG-132-treated MIN6 cell lysates among the KIF5B KD system against STIP1. Note that the Hsp90 level in the STIP1 immunoprecipitants (IP) was greatly decreased by KIF5B deficiency. Repeated twice. (F) Schematic representation of the working hypothesis on differential KIF5B- and heat-shock-protein (Hsp)-dependencies of opposing ER clients Ca_V_1.2 and K_ir_6.2 in control (CT) and KIF5B conditional knockout (cKO) mouse beta cells. In cKO cells, Ca_V_1.2 fails in chaperone exchange to undergo ERAD-mediated degradation, but K_ir_6.2 is intact because it is independent on the KIF5B–Hsp machinery. (G and H) Ca_V_1.2 immunocytochemistry of MIN6 cells that had been transduced with EYFP-KIF5B and/or TagRFP-Hsc70 or without them (NT; G); accompanied by their quantification (H). Scale bar, 5 μm. ns, *p* > 0.05; \*\**p <* 0.01, one-way ANOVA, n = 5–13. Arrow in G, enhanced Ca_V_1.2 expression according to dual overexpression. (I) Vesicle IP of non-transduced (NT) and KIF5B- and Hsc70-overexpressing (K5+H70 OE) MIN6 cell lysates against Ca_V_1.2. Asterisks, tagged protein bands. The tagRFP-Hsc70 band was overlapped with a band of possibly ubiquitinated form. Reproduced twice.

We then compared the chaperone- and co-chaperone-binding capacities of Ca_V_1.2 and K_ir_6.2 by vesicle immunoprecipitation (IP) among the KIF5B-KD system (Fig. 7C and D), where the Ca_V_1.2 level was partially rescued by MG-132 treatment. Interestingly, the association of Hsp70 with Ca_V_1.2 tended to be unaltered. In contrast, the association of Ca_V_1.2 with STIP1, Hsp90, and ER-resident chaperones calnexin-1 and derlin-1 decreased significantly. On the other hand, the Hsp70-binding fraction of K_ir_6.2 was unaltered, but K_ir_6.2 barely bound to Ca_V_1.2, calnexin-1, derlin-1, STIP1, or Hsp90. K_ir_6.2 was still co-associated with KIF5B possibly in the context of endosome/lysosome trafficking (Tanaka *et al*., 1998).

Furthermore, we compared the level of interaction between STIP1 and Hsp90 among the KIF5B-KD system in the presence of MG-132 (Fig. 7E). According to vesicle IP, HSP90 binding capacity of STIP1 was reproducibly decreased upon KIF5B deficiency, this suggested that KIF5B mediates STIP1–Hsp90 binding for maintaining the integrity of chaperone exchange machinery. According to these results, we could define a cochaperone property in KIF5B, dedicated to a chaperone exchange machinery for a subset of ER protein folding (Fig. 7F).

To investigate the possibility of the KIF5B–Hsp machinery to augment the beta cell function, we conducted overexpression trials in MIN6 cells to test if Ca_V_1.2 immunofluorescence was upregulated (Fig. 7G and H). Although overexpression of EYFP-KIF5B alone did not apparently affect the Ca_V_1.2 expression, co-overexpression of EYFP-KIF5B and tagRFP-Hsc70 significantly enhanced it. This may suggest a synergistic role of KIF5B–Hsp machinery in the maturation of their clients.

This KIF5B- and Hsc70-overexpression was found to alter the chaperone-binding capacities of Ca_V_1.2 by vesicle IP. The binding capacities of Ca_V_1.2 to calnexin-1 and Hsp90 were predominantly increased, but that to derlin-1 was decreased (Fig. 7I). The increase in Hsp90-bound Ca_V_1.2 nicely supported our hypothesis that KIF5B enhances the Hsp70-to-Hsp90 chaperone exchange. Calnexin-1 serves for efficient protein production through the calnexin/calreticulin cycle (Kozlov & Gehring, 2020), and its gene silencing significantly increased the Ca_V_1.2 production rate in mouse neonatal cardiomyocytes (Bousette *et al*., 2014). Thus, overproduced Ca_V_1.2 protein may be partially accumulated in this rate-limiting step before exiting the ER. As derlin-1 behaves like a protein degrading chaperone for ERAD (Altier *et al*., 2011), the decrease in the Ca_V_1.2–derlin-1 complex was also reasonable. Accordingly, KIF5B–Hsp system appeared to enhance the Ca_V_1.2 protein folding via the ER chaperones.

### KIF5B facilitates Hsp90-containing microdroplet dynamics on ER sheets

To visualize the KIF5B–Hsp machinery in living cells, we co-transduced primary cKO beta cells with Hsp90-tagRFP and KIF5B-EYFP and conducted live cell imaging. As revealed by stereoscopic presentation of *z-*stack images, tagRFP-Hsp90 and KIF5B-EYFP tended to co-accumulate on patch-like microdroplets especially in the cell bottom (Fig. EV2E). Time-lapse microscopy revealed that Hsp90 formed fine meshwork, along which the microdroplets containing Hsp90 and KIF5B underwent very dynamic movements (Fig. 8A and Movie EV5). Occasionally, KIF5B- and Hsp90-containing tubules were extended from one microdroplet toward another, suggesting their dynamic material-exchanging properties that may serve for liquid-liquid phase separation (LLPS) (Naz *et al*., 2024).

**Figure 8.**
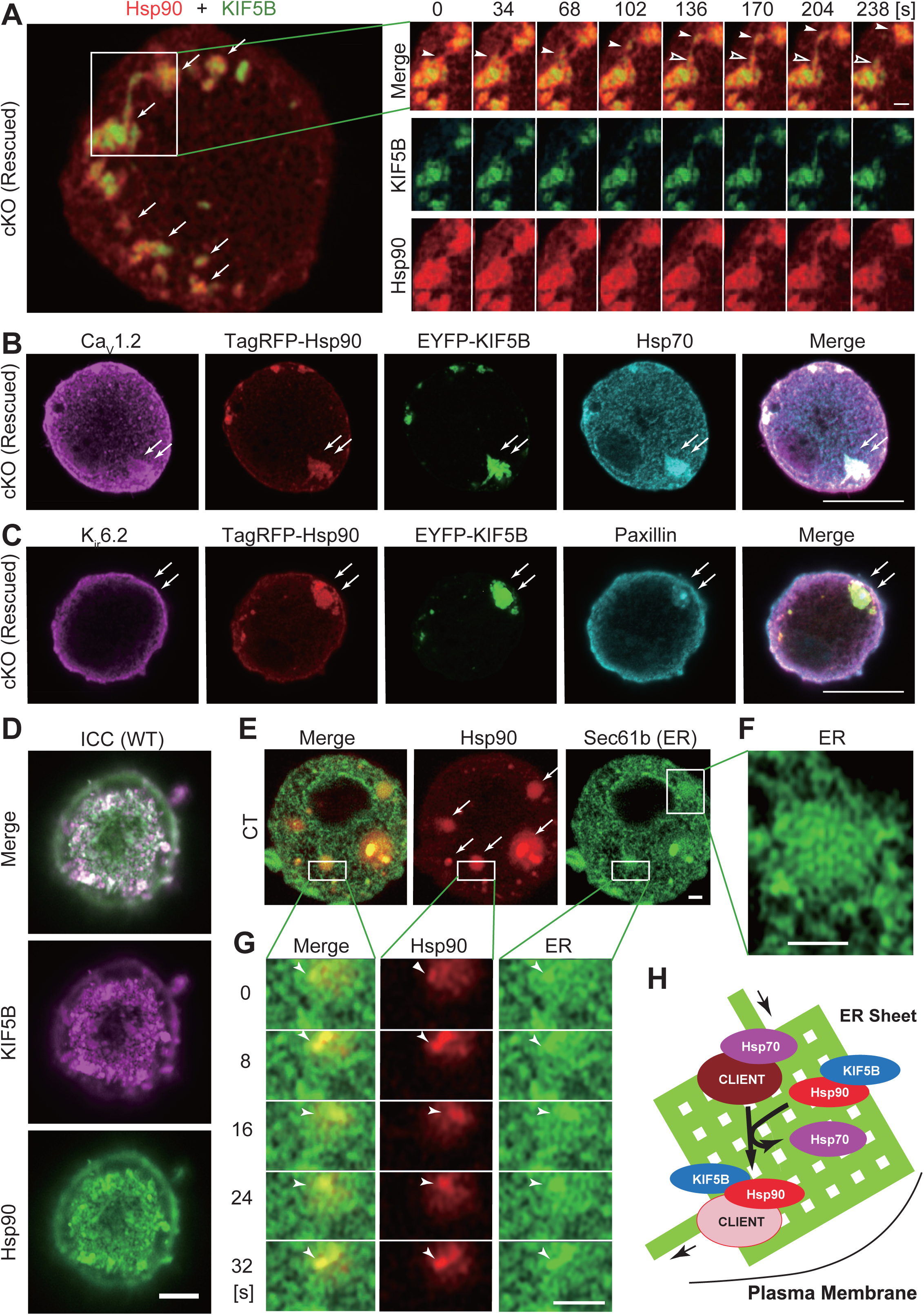
KIF5B recruits Hsp90 onto ER sheets for proper Ca_V_1.2 protein folding. (A) Time lapse imaging in the bottom of a rescued primary cKO beta cell, expressing tagRFP-Hsp90 (red) and KIF5B-EYFP (green). Arrows, microdroplets. Note that two co-accumulated microdroplets appeared to exchange their components through a double-labeled dynamic tubule (open and closed arrowheads). Corresponding to Fig. EV2E and Movie EV5. (B and C) Immunofluorescence microscopy in the bottom of primary cKO beta cells expressing tagRFP-Hsp90 (red) and KIF5B-EYFP (green), against Ca_V_1.2 and Hsp70 (B) and against K_ir_6.2 and Paxillin (C). Scale bars, 10 μm. Arrows, co-accumulated microdroplets. Reproduced 5–10 times. (D) Immunocytochemistry of a wild type islet cell against KIF5B and Hsp90 using Airyscan microscopy. Reproduced twice. (E–G) Live fluorescence microscopy of the bottom of CT primary beta cells expressing tagRFP-Hsp90 (red) and the ER marker mEmerald-Sec61β (green) using Airyscan microscopy with low (E) and high (F, G) magnifications and in a time sequence (G). Scale bars, 1 μm. Arrows, microdroplets. Arrowheads, a portion of the ER sheet accompanied by Hsp90 microdroplets. Corresponding to Movie EV6. (H) Schematic representation of a ER sheet where the Hsp70-to-Hsp90 chaperone exchange occurs for the quality control of ER client protein folding by the help of KIF5B molecular motor.

According to immunocytochemistry of KIF5B-rescued cKO beta cells, Hsp70, Ca_V_1.2, and the focal adhesion marker paxillin tended to co-accumulate on the microdroplets with KIF5B and Hsp90 (Fig. 8B and C). However, K_ir_6.2 was excluded from there, reflecting its independence on Hsp90. The colocalizing features between KIF5B and Hsp90 on microdroplets were further confirmed by intrinsic immunocytochemistry of wild type primary beta cells (Fig. 8D).

To investigate the relationship between the ER and those microdroplets, we co-transduced primary beta cells with tagRFP-Hsp90 and the ER marker, mEmerald-Sec61β (Nixon-Abell et al, 2016)(Fig. 8E–G). Airyscan microscopy revealed that Hsp90-including microdroplets consisted of a bright core and rather amorphous halo. They were frequently colocalizing to finer parts of ER meshwork (Fig. 8F), which were termed “ER sheets” (Shibata *et al*., 2010). Time-lapse analysis further revealed that those ER sheets and Hsp90 dynamically migrated together (Fig. 8G and Movie EV6).

These data collectively suggested that KIF5B is essentially involved in Hsp70-to-Hsp90 ER chaperone exchange system that occurs in microdroplets associated with ER sheets, and this may facilitate the proper maturation of a subset of ER clients including Ca_V_1.2 (Fig. 8H). This may be the subcellular basis of KIF5B-mediated control of Ca_V_1.2 expression that may finely tune the GSIS for maintaining glucose homeostasis.

## Discussion

In the present study, we found that KIF5B kinesin is essential for the first- and second-phase GSIS, mainly through a protein folding process in the ER sheets involving the KIF5B- and Hsp90-containing microdroplets (Fig. 8). These microdroplets may be transported and scaffolded by the KIF5B molecular motor, and may be essential for Hsp70-to-Hsp90 chaperone exchange as well as ER chaperone binding for proper folding of a subset of membrane proteins including Ca_V_1.2 (Figs. 6 and 7). Ca_V_1.2 enterprises glucose-stimulated Ca^2+^ transients that controls first- and second-phase GSIS, partly through Ca^2+^-dependent SNARE complex formation (Trexler & Taraska, 2017) and partly through cortical F-actin remodeling that modulates insulin granule dynamics (Kalwat & Thurmond, 2013). This finding uncovers an outstanding molecular mechanism in the stimulation-secretion coupling of GSIS that acts against diabetes, as well as the relevance of kinesin-1 molecular motor in protein folding in the ER. Because our beta-cell specific KIF5B cKO mice largely phenocopied the Ca_V_1.2 cKO mice (Schulla *et al*., 2003), rather than Ca_V_2.3 KO mice (Jing *et al*., 2005), and that the Ca_V_1.2 agonist Bay-K 8644 nicely restored the GSIS in KIF5B cKO islets (Fig. 5F), the Ca_V_1.2 channel was considered as the most physiologically relevant client of the KIF5B–Hsp machinery in pancreatic beta cells.

The relationship between ER and kinesin-1 has long been suggested by many groups (Gupta *et al*., 2008; Raiborg *et al*., 2015; Woźniak *et al*., 2009), but the KIF5B’s specific role in protein folding has not been determined yet. In our previous knockout mouse study (Tanaka *et al*., 1998), we reported that the overall ER distribution and BFA-mediated Golgi-to-ER trafficking were largely unaltered, but mitochondria and lysosomes revealed significant perinuclear clustering in KIF5B-deficient extraembryonic cells. On the other hand, Allan and colleagues showed that dominant-negative kinesin-1 inhibits centripetal ER tubule motility in VERO cells (Woźniak *et al*., 2009), suggesting that kinesin-1 still serves as one of the molecular motors that synergistically act on ER morphogenesis. As of associating factors, the Rab18 and kinectin-1 (KNT-1) interaction may facilitate the dynamics of ER-focal adhesion (FA) contact as well as kinesin-1-dependent ER extension (Guadagno *et al*., 2020; Zheng *et al*., 2022), and KNT-1 is reported to be enriched in ER sheets (Shibata *et al*., 2010). Recently, it has also been reported that KIF5B transports vimentin (Robert *et al*., 2019), and that vimentin controls ER remodeling during ER stress (Cremer *et al*., 2023). These findings support our current idea that KIF5B regulates the structure and function of the ER in beta cells. As we have shown in an overexpression study (Fig. 7G–I), these KIF5B- and Hsp-dependent processes enhance the production of Ca_V_1.2 protein in pancreatic beta cells, to serve for maintaining the stimulation-secretion coupling for glucose homeostasis.

Our data suggested that KIF5B deficiency augmented the basal insulin secretion as well as constitutive depolarization (Figs. 1F and 4I). This could be mostly explained by the downregulation of Na/K ATPase and BK_Ca_ in KIF5B KD cells (Fig. 5A and B). Alternatively, Ca_V_1.2 knockout mice exhibited a similar elevation of basal insulin secretion with significant GSIS decrease (Schulla *et al*., 2003). Because Ca_V_1.2 augments BK_Ca_ activity through a direct interaction (Plante *et al*., 2021), the Ca_V_1.2 decrease itself may also account for the baseline changes in the KIF5B cKO cells.

Regarding the second-phase insulin secretion, our data have revealed that KIF5B is essential for a glucose-stimulated sequence of SFK and Rho-family GTPase activation (Fig. 4E–G), actin remodeling (Fig. 4A and B), and changes in insulin granule kinetics and full-fusion exocytosis probability (Figs. 2 and 3), which collectively maintain the second-phase GSIS. Although the pSFK–Cdc42–PAK1–Rac1 signaling pathway can partly crosstalk with Ca_V_1.2-mediated Ca^2+^ transients (Veluthakal & Thurmond, 2021), the glucose-stimulated activation mechanism of SFK within 1 min is still elusive as far as we know. The significant decrease of PIP_2_ in KIF5B-deficient beta cells (Fig. 5G and H) may affect PIP_2_-dependent FAK activation (Zhou *et al*., 2015) as a prerequisite to FAK–SFK coactivation (Huveneers & Danen, 2009). We have previously identified that KIF26A and KIF21B kinesins regulate the FAK–SFK interaction and Rac1 nucleotide cycling, respectively, in neurons (Morikawa *et al*., 2018; Wang *et al*., 2018). If our prediction was correct, gene manipulation of those related kinesins, as well as that of KIF5B (Fig. 7G and H), can enhance GSIS, which will be subjected to future research.

In KIF5B cKO beta cells, a small first-phase peak of GSIS was still remaining (Fig. 1F). Because Ca^2+^ transients and pSFK elevation were almost completely abolished (Fig. 1K and L), this peak should be driven by other second messenger systems possibly involving protein kinases A and C (Komatsu *et al*., 2013) and/or Na^+^ channels (Shigeto *et al*., 2006). KIF5B-deficient beta cells will thus serve as an ideal system for studying a Ca^2+^- or SFK-independent component of first-phase GSIS, which will contribute to next-generation diabetes therapeutics (Sola *et al*., 2015; Wang *et al*., 2021).

In this study, we investigated the role of KIF5B in ERQC system for the Ca_V_1.2 protein, which may be especially relevant for proper GSIS in beta cells. These cell biological findings will provide insights into the role of kinesin-1 in insulin secretion, from a completely unexpected view of cell biology. These findings will greatly stimulate future research and development on islet cell biology. Simultaneously, they will progress the basic cell biology of ER compartmentalization, which will be mostly dedicated to the geographical understanding of essential subcellular mechanisms.

## ACKNOWLEDGMENTS

The authors thank Tetsuo Noda (Cancer Inst, Japan) for ES cell technology, Mitsuhisa Komatsu (Shinshu Univ) for islet recovery, Junqing Sun (Univ Tokyo) for patch clamp recording, Phil Soriano (Mt. Sinai School of Medicine) for the *ROSA-STOP* mice, Michael Sixt and Roland Wedlich-Soldner (Max Plank Institute of Biochemistry, Germany) for the *Lifeact-EGFP* mice, Claes B. Wollheim (University Medical Center, Geneva, Switzerland) for the Ins1 cells, Jun-ichi Miyazaki (Osaka University) for the MIN6 cells, Jonathan Jones (Northwestern University Medical School) for the 804G cells, Gero Miesenboeck (Sloan-Kettering Institute for Cancer Research) for the *synapto.pHluorin* cDNA, Jennifer Lippincott-Schwartz (NIH) for the *mEmerald-Sec61β* expression vector, Michiyuki Matsuda (Kyoto Univ) for the *Raichu* biosensor cDNAs, and Luis F. Santana (University of Washington) for the *Ca_V_1.2-EGFP* cDNA. We are also grateful to Yuki Hashimotodani (Doshisha Univ), Takuya Okada (Physio-tech, Co., Ltd., Japan), Ingo Kleppe and Bernhardt Zimmermann (ZEISS), Fumiyoshi Ishidate (Kyoto Univ); and Shuo Wang, Yoshinobu Shimazawa, Tatemitsu Rai, Shinsuke Niwa, Kyosuke Nakajima, Shuzo Hasegawa, Momo Morikawa, Nobuhisa Onouchi, Takeshi Akamatsu, Hiromi Sato, Haruyo Fukuda, and previous members of the Hirokawa laboratory for technical assistance, support, and valuable discussions. This study was supported by JSPS KAKENHI grant numbers JP23000013 and JP16H06372 to N.H. and JP20K06634 to Y.T.; a research grant-in-aid from JEOL Ltd. to N.H.; Research Grants-in-Aid from Uehara Memorial Foundation, ONO Medical Research Foundation, and Japan IDDM Network to Y.T.

## AUTHOR CONTRIBUTION

Project administration and conceptualization by NH; Conceptualization and methodology by YT and MN; Investigation by YT, AF, WY, HU, and MN; Funding acquisition by NH and YT; Original writing by YT; Review and editing by NH, MN, and YT.

## Materials and Methods

### Mouse models

For generating KIF5B cKO mice, homologous recombination in the ES cells was conducted. A promoter-trap 3-*loxP* type targeting vector floxing the 74 bp p-loop exon (Fig. EV1A) was transfected into the J1 line of mouse embryonic stem (ES) cells by electroporation, as previously described (Tanaka *et al*., 1998). Homologous recombinants were screened by genomic Southern blotting with an efficiency of more than 70% (Fig. EV1B). Then, the *pCre-Pac* vector was transiently transfected into the homologous recombinants in the presence of puromycin, to generate *2-loxP* alleles as previously described (Teng *et al*., 2005). Each colony was genotyped with PCR using the following primer sets: neo, 5’-TGGGCACAACAGACAATCGG-3’ and 5’-ACTTCGCCCAATAGCAGCCAG-3’; 5’-floxed region, 5’-CCAGATAACAGTTAAAAGCAGTGAAGG-3’ and 5’-CCATTATAGCCCTCAAGAACATCTATG-3’; 3’-floxed region (flox), 5’-CCCACACGATGGAGGTAATGTTTC-3’ and 5’-CCTGGCTGATATAGACAATCTTATGAGAAG-3’; and Cre, 5’-AGGTTCGTTCACTCATGGA-3’ and 5’-TCGACCAGTTTAGTTACCC-3’ (Fig. EV1C). The *2loxP* allele was transmitted to the germline using blastocyst injection to establish the line #38. Rat insulin promoter-driven *Cre* (*Rip2-Cre*) transgenic mice (Kulkarni *et al*., 1999; Postic *et al*., 1999) were obtained from the Jackson Laboratory and crossed with line #38. The ROSA-STOP reporter mice (Soriano, 1999) were kindly provided by Dr. P. Soriano (Mt. Sinai School of Medicine) and crossed with *Rip2-Cre* mice to verify the specificity of Cre-expressing cells (Fig. EV1D). KIF5B deficiency in cKO cells was verified by immunofluorescence microscopy as described previously (Tanaka *et al*., 1998). These mice were maintained on a C57BL/6J background in a specific pathogen-free environment under a 14/10-h light/dark cycle under the institutional regulations. *Lifeact-mCherry* mice were a kind gift of Drs. Michael Sixt and Roland Wedlich-Soldner (Max Planck Institute for Biochemistry, Germany) (Riedl *et al*., 2010).

Institutional approval for mouse experiments was received from the Institutional Animal Care and Use Committee of the University of Tokyo Graduate School of Medicine (#MP15-118, #MP20-92).

### Histology

For immunohistochemistry and enzyme histochemistry of the tail region of the pancreas, mice were anesthetized and perfused with 4% paraformaldehyde in PBS to generate cryosections. β-galactosidase activity was stained with 0.5 mg/mL X-gal, 10 mM K_3_[Fe(CN)_6_], 10 mM K_4_[Fe(CN)_6_], 0.01% Na desoxycholate, 0.02% Nonidet P-40, 20 mM Tris-HCl, and 100 mM K-phosphate buffer [pH 7.4] according to previously described methods (Joyner, 2000). Immunohistochemistry was performed as previously described (Kanai *et al*., 2000). Fixed cells were labeled with a rabbit anti-KIF5B antibody (Tanaka *et al*., 1998) and a mouse anti-insulin antibody (Sigma), followed by Alexa-conjugated secondary antibodies (Invitrogen), observed using LSM510 or LSM710 confocal laser scanning microscopes (ZEISS), and analyzed using the MosaicJ plugin (Thevenaz & Unser, 2007) on ImageJ software (Abramoff *et al*., 2004).

### Blood sugar test

The intraperitoneal glucose tolerance test (IPGTT) was performed as previously described (Ohtsubo *et al*., 2005; Yang *et al*., 2014). In brief, 4-month-old male mice were starved overnight and intraperitoneally injected with 2 g glucose/kg body weight. Their blood glucose levels were measured at the indicated time points using a Medisafe-Mini blood glucose meter (Terumo, Tokyo, Japan).

### Pancreatic islets

Pancreatic islets were recovered from male mice using 8 g/L collagenase digestion (C-7657, Sigma) in Krebs Ringer Bicarbonate (KRB) solution (Daniel *et al*., 2002) as described previously (Szot *et al*., 2007; Yang *et al*., 2014). Prior to the experiments, the islets were cultured in RPMI-1640 medium (Invitrogen) with 10% FCS and antibiotics in a 5% CO_2_ atmosphere at 37°C for 3 h to overnight as described previously (Carter *et al*., 2009).

Islet perifusion assay (Fig. 1F) was performed as previously described (Noda *et al*., 1996). 6–8-month-old mouse islets were stimulated by KRB solution containing 20 mM glucose, following a preincubation with KRB solution containing 2 mM glucose. The perifusates were subjected to ELISAs using LBIS Mouse Insulin ELISA Kit (WAKO-Shibayagi, Japan) according to the manufacturer’s protocol. Only the samples that reverted to the initial level of secretion after the stimulation period were subjected to the statistical analyses. The mean insulin secretion on the period without stimulants was calculated as basal insulin secretion of each trial. Increments in 0–10 min and that in 10– 44 min over the basal level secretion were calculated as first- and second-phase GSIS, respectively (Fig. 1G).

For the bulk insulin secretion assay (Fig. 5F), four islets from adult cKO mice in each tube were preincubated in KRB solution supplemented with 2.8 mM glucose. Then they were once rinsed with the same solution, and stimulated with 1 mL of KRB solution containing either 2.8 mM glucose, 16.7 mM glucose, or 16.7 mM glucose plus 1 μM Bay-K 8644 (#S7924, Selleck) at 37°C for 1 h in a water bath. The supernatants were subjected to LBIS Mouse Insulin ELISA Kit (Luminescent type; WAKO-Shibayagi, Japan) according to the manufacturer’s protocol.

The ATP/ADP ratio was measured as described previously (Lu *et al*., 2010) using an ApoSENSOR ADP/ATP Ratio Assay Kit (#K255-200, BioVision) following the manufacturer’s instructions.

### Primary culture of pancreatic beta cells

For the primary culture, Lab-Tek chambered coverslips (Nalge Nunc) or 35 mm glass bottom dishes (Matsunami) were coated with a conditioned medium of 804G cells (Bosco *et al*., 2000; Langhofer *et al*., 1993). Precultured islets were incubated with 0.04% EDTA in PBS at 37°C for 10 min and gently dissociated into single cells by pipetting. Following a medium change through centrifugation, islet cells were plated and cultured in KRB medium or in Ins1 medium [10 mM HEPES pH 7.4 (Gibco), 10% fetal bovine serum (Sigma Aldrich), 1 mM Na pyruvate (Gibco), 50 mM 2-mercaptoethanol (Sigma Aldrich), penicillin-streptomycin (Gibco), RPMI1640 with L-glutamine (Gibco)] (Asfari *et al*., 1992) in a 5% CO_2_ atmosphere at 37°C and subjected to analyses within 1 week. Beta cells were identified as round and > 10 *μ*m cells. Coating with 804G conditioned medium significantly stabilized the attachment of beta cells to the matrix and improved their viability.

### Cell lines

The MIN6 cells (Miyazaki *et al*., 1990) were kindly provided by Dr. Jun-ichi Miyazaki (Osaka University). The 804G cells (Bosco *et al*., 2000) were kindly provided by Dr. Jonathan Jones (Northwestern University Medical School). The Ins1 cells (Asfari *et al*., 1992) were kindly provided by Dr. Claes B. Wollheim (University Medical Center, Geneva, Switzerland).

### Antibodies

A rabbit anti-Ca_V_1.2 antibody (N-17-R, #sc-16229-R, RRID:AB_2228387), a rabbit anti-K_ir_6.2 antibody (H-55; #sc-20809; RRID:AB_2130466), a goat anti-PIP5Kα (PIPKIα) antibody (M-20, #sc-11775; RRID:AB_2268303), and a goat anti-SUR1 antibody (N-18, #sc-11226; RRID:AB_2130475) were purchased from Santa Cruz Biotechnology; a mouse anti-PIP_2_ IgM antibody (#Z-A045, RRID:AB_427211) was from Echelon Research labs; a rabbit anti-GFP antibody (#598, RRID:AB_591819) was from MBL; a mouse anti-LC3 antibody (Clone LC3-1703, #CTB-LC3-2-IC, RRID:AB_10707197) was from Cosmo Bio; a rabbit anti-BK_Ca_ (K_Ca_1.1) antibody (#APC-151, RRID:AB_10915895) and a rabbit anti-Ca_V_2.3 antibody (#ACC-006, RRID:AB_2039777) were from Alomone Labs; a mouse anti-syntaxin-1 antibody (#MAB336, RRID:AB_2196527) was from Millipore; a mouse anti-Na/K ATPase beta 2 antibody (#610914; RRID:AB_398231) and a mouse anti-paxillin antibody (#610051, RRID:AB_397463) were from BD Transduction Labs; a mouse anti-Hsc70/Hsp70 antibody (Clone BB70, #ADI-SPA-822-D, RRID:AB_2039252) and a rat anti-Hsp90 antibody (Clone 16F1, #ADI-SPA-835-D, RRID:AB_2039281) were from Enzo; a rabbit anti-STIP1 antibody (#15218-1-AP; RRID:AB_2255518) and a rabbit anti-calnexin-1 (CANX) antibody (#10427-2-AP, RRID:AB_2069033) were from Proteintech; a mouse anti-derlin-1 antibody (#SAB4200148, RRID:AB_10624068), an anti-α-tubulin antibody (Clone DM1A; #CP06; RRID:AB_2617116), and an anti-insulin antibody (Clone K36aC10, #I-2018, Sigma Aldrich; RRID:AB_260137) were from Sigma Aldrich; a rabbit anti-phospho Src antibody for pSFK (#ab32078; RRID:AB_2286707) was from Epitomics/Abcam; and a rabbit anti-KIF5B antibody (RRID:AB_2571745) was previously described (Tanaka *et al*., 1998). Alkaline-phosphatase-conjugated anti-rabbit IgG, anti-chick IgG and anti-goat IgG antibodies were purchased from Cappel. Alexa-conjugated phalloidin, anti-rabbit IgG, anti-mouse IgG and anti-goat IgG antibodies were purchased from Thermo Fisher.

### Gene silencing

For gene silencing, antisense miRNAs were transcribed from mammalian PolII-based expression vectors derived from a Block-iT Pol II miR RNAi Expression vector kit (Thermo Fisher). For *Kif5b* knockdown, the oligonucleotides 5’-TGCTGTTTCAGGGCTGACTCCAAAGCGTTTTGGCCACTGACTGACGCTTTGGACAGCCCTGAAA-3’ and 5’-CCTGTTTCAGGGCTGTCCAAAGCGTCAGTCAGTGGCCAAAACGCTTTGGAGTCAGCCCTGAAAC-3’ (KD1); and 5’-TGCTGTTCTCTGTGACTCTGGATCTGGTTTTGGCCACTGACTGACCAGATCCAGTCACAGAGAA-3’ and 5’-CCTGTTCTCTGTGACTGGATCTGGTCAGTCAGTGGCCAAAACCAGATCCAGAGTCACAGAGAAC-3’ (KD2) were obtained from Invitrogen, annealed, and inserted into the provided vector with *tagRFP* cDNA as an expression marker and equally sujected to the knockdown analyses to obtain consistent results. For *Stip1* knockdown, the oligonucleotides 5’-CCTGTGCAGCCGTCCATAGGGCCTGTCAGTCAGTGGCCAAAACAGGCCTATGAGGACGGCTGCAC-3’ and 5’-TGCTGTGCAGCCGTCCTCATAGGCCTGTTTTGGCCACTGACTGACAGGCCTATGGACGGCTGCA-3’ (#Mmi520981, Invitrogen) were used. As a negative control, a scrambled sequence provided by the kit were used to exclude the possibility of nonspecific adverse effects of miRNA expression. Those miRNA expression vectors were introduced into adenoviral vectors using a ViraPower Adenoviral Expression System, purified by CsCl centrifugation, and subjected to cells according to manufacturer’s protocols.

### Expression vectors

For generation of a *TagRFP-Hsc70* expression vector, the *Hsc70* cDNA was recovered from a previously described *ECFP-Hsc70* expression vector (Yang *et al*., 2014) by *SalI* and *XhoI,* enzymes, and ligated to *TagRFP-C* vector (Evrogen).

For generation of a *TagRFP-Hsp90* expression vector, the FANTOM I1C0020P08 clone (RIKEN) containing *Hsp90aa1* cDNA (MGI:3564069) was amplified by 5’-ACCCTCGAGCTATGCCTGAGGAAACCCAGACCCAAG-3’ and 5’-ACCGGTACCTTAGTCTACTTCTTCCATGCGTGATGTGTC-3’ and ligated into *pTagRFP-C vector* (Evrogen) with *Xho*I and *Kpn*I sites.

For the *phogrin*-*Dronpa-Green1* and *phogrin-EGFP* expression vectors, mouse *phogrin* cDNA was ligated with *pDronpa-Green1* (MBL, Japan) or *pEGFP-N1* (Clontech), and transferred to the ViraPower Adenoviral Expression System (Thermo Fisher).

The *mEmerald-Sec61β* expression vector was a kind gift from Prof. Jennifer Lippincott-Schwartz through Addgene #90992 (Nixon-Abell *et al*., 2016). The *Ca_V_1.2-EGFP* expression vector was a kind gift from Dr. Luis F. Santana (Navedo *et al*., 2010). A *synapto.pHluorin* Adenoviral expression vector was kindly provided by Dr. Gero Miesenboeck, Sloan-Kettering Institute for Cancer Research (Miesenbock *et al*., 1998). The FRET biosensors *pRaichu 1011x* and *pRaichu 1054x* were kind gifts of Dr. Michiyuki Matsuda (Univ Kyoto).

For rescuing the KIF5B cKO cells, mouse *Kif5b* full-length cDNA (Tanaka *et al*., 1998) was ligated into the *pEYFP-N1, pEYFP-C1* or *pECFP-C1* vectors (Clontech). The expression vectors were transduced into primary islet cells using the ViraPower Adenoviral Expression System (Thermo Fisher) or using lipofectamine-LTX transfection reagent (Thermo Fisher). The mobility of expressed proteins on SDS–PAGE was verified by immunoblotting of transfected rat Ins1 cell lysates (Asfari *et al*., 1992). For KIF5B- and Hsc70-overexpression, MIN6 cells were transduced by Adenoviral expression vectors for ECFP-KIF5B and/or tagRFP-Hsc70 for 48 h, and subjected to immunocytochemistry or vesicle IP.

### Pharmacology

Cytochalasin B (CyB) was purchased from Sigma Aldrich (#C6762) and applied to cells at 10 μg/mL (Lacy *et al*., 1973). Ionomycin was purchased from Sigma Aldrich (#I9657) and applied to cells at 1 μM (Sekine *et al*., 2006). For the VGCC antagonists, 20 μM nifedipine (#N7634, Sigma Aldrich) against L-type VGCC (Gilon *et al*., 1997), 1 μM SNX482 (#RTS-500, Alomone Labs) against R-type VGCC (Bourinet *et al*., 2001), and 0.3 mM L-ascorbate (#A4034, Sigma Aldrich) against T-type VGCC (Nelson *et al*., 2007) were applied to cells for a negative control for electrophysiology. Bay-K 8644 was purchased from Selleck (#S7924) and applied to islets at 1 *μ*M (Satin *et al*., 1995). Cycloheximide was purchased from Fuji-Film WAKO (#037-20991) and applied to cells at 30 *μ*g/mL (Yang *et al*., 2014), in accordance with 25 μM leupeptin (#336-40413, Fuji-Film WAKO) or 10 μM MG-132 (#P1102, Enzo) as described (Chou & Deshaies, 2011; Di Biase *et al*., 2011).

### Fluorescence microscopy

Immunofluorescence microscopy was performed as described previously (Yang *et al*., 2014) using CanGetSignal reagents according to the manufacturer’s protocol (Toyobo). Especially, primary cultured mouse pancreatic beta cells were fixed with 2% paraformaldehyde (PFA)/0.1% glutaraldehyde (GA) at 37°C for 10 min, permeabilized with 0.1% Triton X-100 in PBS for 5 min at room temperature, blocked in 10% bovine serum albumin (BSA) in PBS for 10 min at room temperature, and subjected for immunostaining using Can Get Signal Immunostain Solution A (TOYOBO) according to the manufacturer’s protocol. The samples were then subjected to a confocal laser-scanning microscope (LSM710 equipped with a GaAsP detector and LSM780-Airyscan, ZEISS), fast confocal laser-scanning microscope (LSM 5LIVE-Duo, ZEISS), or a TIRF/STORM microscope (ELYRA P.1, ZEISS, equipped with an iXon+ EM-CCD camera, Andor) as previously described (Nakata *et al*., 2011; Tanaka *et al*., 2016; Wang *et al*., 2022) and analyzed using ImageJ software (Schindelin *et al*., 2015). For quantification of the membrane channels in an optical section, an area within 0.8 μm of the surface was selected and measured for mean signal intensity.

For degradation assays, cells were treated with 30 μg/mL cycloheximide (CHX) with either 25 μM leupeptin (Sigma Aldrich) or 10 μM MG-132 (Merck) as described (Chou & Deshaies, 2011; Di Biase *et al*., 2011).

For the BFA washout assay, the cells were transduced with a rabbit Ca_v_1.2-EGFP expression vector (Navedo *et al*., 2010) in the presence of 1 μM brefeldin A (Sigma Aldrich) at 19.5°C overnight. Then, the cells were washed out using complete medium twice, cultured at 37°C for the indicated times, and subjected to confocal laser scanning microscopy as described previously (Nakata & Hirokawa, 2003; Tanaka *et al*., 2023).

Insulin granule exocytosis was imaged using TIRF microscopy with a *synapto.pHluorin* adenoviral expression vector (Miesenbock *et al*., 1998; Tsuboi & Rutter, 2003), treated with secretagogues and/or drugs as indicated, and observed with an ELYRA P.1 microscope (ZEISS) at 30 ms intervals for 1 min, as previously described (Aoki *et al*., 2010; Ohara-Imaizumi *et al*., 2007). For F-actin imaging, cells were treated with 10 μg/mL cytochalasin B (CyB; Sigma Aldrich) (Lacy *et al*., 1973), or 1 μM ionomycin (Sigma Aldrich) (Sekine *et al*., 2006), fixed, and stained with Alexa-conjugated phalloidin (Invitrogen) according to the manufacturer’s protocols. For time-lapse analysis of F-actin, primary beta cells were recovered from *Lifeact-EGFP* transgenic mice, which were kindly provided by Drs. Michael Sixt and Roland Wedlich-Soldner (Max Planck Institute for Biochemistry, Germany) (Riedl *et al*., 2010), and subjected to *Kif5b* gene silencing and observed in time-lapse analysis with an LSM-710 microscope (ZEISS).

For observation of the cortical insulin granules, the *phogrin-Dronpa-Green1* cDNA expression vector was transduced to primary beta cells using ViraPower adenovirus system (Invitrogen). The cells were then fixed in 4% paraformaldehyde/PBS for 10 min and observed using a TIRF/PALM microscope (ZEISS). For tracking of granule movements, primary beta cells were transduced with phogrin cDNA ligated with *pEGFP-N1* vector (Clontech), treated successively with Ins1 media containing 2 mM and 20 mM glucose respectively for 30 min, and observed using an LSM 5LIVE-Duo microscope (ZEISS; 30 frames/17.7 s) or a CSU-X1 spinning disc microscope (Yokokawa; 200 frames/100 s) immediately after removing the cells from the CO_2_ incubator. The images were analyzed using Imaris software for particle tracking (Bitplane AG).

### Imaging of F-actin

For F-actin imaging, the cells were fixed using half-Karnovsky fixative [2% paraformaldehyde, 2.5% glutaraldehyde, and 0.1 M cacodylate buffer (pH 7.4)] at 37°C for 15 min, permeabilized with 0.1% Triton X-100 in PBS at room temperature for 5 min, blocked with 10% BSA in PBS at room temperature for 10 min, stained with phalloidin conjugated with 4–10 U/mL of Alexa-568 or Alexa-488 (Invitrogen) in the blocking solution, washed with PBS for 5 min three times, observed using a confocal laser scanning microscope (LSM 5LIVE-Duo, LSM710, LSM780-Airyscan, ZEISS).

For quantification of cortical F-actin (Fig. 4), the cortical fluorescent signals of optical sections, approximately at the level of half of the cell height, were selected with the help of the “Cell Outliner” plugin of the ImageJ software, and the median signal of each selection was analyzed.

### Rho-family GTPase activities

For imaging the activities of Rac1 and Cdc42 GTPases, *pRaichu* 1011x and 1054x vectors (Kiyokawa *et al*., 2006), kindly provided by Dr. Michiyuki Matsuda, Kyoto University, were transduced into the cells via adenoviral vectors, respectively. Cells were then pretreated with Ins1 medium containing 2 mM glucose for 30 min and stimulated with 20 mM glucose for the indicated period. The YFP/CFP ratio was then measured at 458 nm excitation using an LSM 5LIVE-Duo confocal microscope (ZEISS) and quantified using ImageJ software (Abramoff *et al*., 2004).

### Ca imaging

Ca^2+^ imaging was performed with loading the cell with Fluo4-AM (Dojindo, Japan) as previously described (Tanaka *et al*., 2016; Tanaka *et al*., 2005; Wang *et al*., 2018), using LSM710 and LSM780 confocal laser scanning microscopes (ZEISS). The recording was started with Ins1 medium without glucose, and then continued with stimulation with 20 mM glucose. Finally, the medium was supplemented with 30 mM CaCl_2_, 125 mM KCl, and 10 μM ionomycin, and then washed with DDW, to normalize the experimental data as ΔF/Fmax.

### Patch clamp recording

Patch clamp recording of primary cultured beta cells at DIV1–2 were performed as previously described (Rorsman & Trube, 1986). The bath solution was 138 mM NaCl, 5.6 mM KCl, 1.2 mM MgCl_2_, 2.6 mM CaCl_2_, and 10 mM HEPES-NaOH [pH 7.4]; with 0 or 10 mM of glucose. The pipette was filled with 125 mM KCl, 30 mM KOH, 4 mM MgCl_2_, 3 mM Na_2_ATP, 2 mM CaCl_2_, 10 mM EGTA, and 5 mM HEPES-KOH [pH 7.15]. The intracellular potential was recorded stepwise under 0 and 10 mM of glucose in the external medium. Inward Ca^2+^ currents were recorded with a voltage clamp under 10 mM glucose with or without VGCC inhibitor cocktail that is 20 *μ*M nifedipine against L-type VGCC (Gilon *et al*., 1997), 1 μM SNX482 against R-type VGCC (Bourinet *et al*., 2001), and 0.3 mM ascorbate against T-type VGCC (Nelson *et al*., 2007). The data acquisition and processing were performed using Axon pCLAMP 10 software (Molecular Devices).

### Immunoprecipitation

Immunoprecipitation was performed as previously described (Ueno *et al*., 2011). MIN6 cells were transduced with scrambled control (SC) or KIF5B knockdown (KD) adenoviral vectors for 6 days. They were then treated by 10 μM MG-132 for 6 h, and harvested using 10 mM HEPES [pH 7.4], 150 mM NaCl, 0.1% Triton-X 100, and protease inhibitors (Roche). Postnuclear fractions were precipitated against protein A Sepharose beads (GE Healthcare) or *μ*MACS Protein A Microbeads (Miltenyi Biotech) conjugated with 2 *μ*g of antibody or normal rabbit IgG (Cappel). After extensive washing, the beads were boiled with 2 × Laemmli’s sample buffer and subjected to immunoblotting using an Amersham ECL Prime Western Blotting Detection System (GE Healthcare).

### Proximity ligation assay

Proximity ligation assay was conducted using a DuoLink system (Sigma Aldrich) with rabbit anti-Ca_V_1.2, mouse anti-Hsp70, and rat anti-Hsp90 antibodies according to the manufacturer’s protocols. The anti-mouse probe of the kit cross-reacted rat IgG, so that an anti-mouse (plus) and anti-rabbit (minus) pair of the Duolink probes was applied. The samples were subjected to a confocal laser scanning microscope (LSM710 and LSM780-Airycan, ZEISS); and *z-*projection at the maximum intensity was conducted using ImageJ ver. 1.54i software.

### qRT–PCR

qRT–PCR was performed as previously described (Tanaka *et al*., 2016; Yang *et al*., 2014). SC/KD-miRNA-transduced MIN6 cells were cultured in 2 mM glucose and treated with a total RNA isolation mini kit (Agilent) and 1^st^ strand synthesis kit (Origine). The 1^st^ strand cDNA was subjected to real-time PCR on a LC480 thermal cycler instrument II (Roche) using SYBR Premix Ex Taq (Tli RNaseH Plus, #RR420, TaKaRa) with the following primers: *K_ir_6.2* cDNA with 5’-CTCATCATCTACCACGTCATCGA-3’ and 5’-GTTTCTACCACGCCTTCCAAGA-3’ (Camerino *et al*., 2013); *Ca_V_1.2* cDNA with 5’-TCCTCATCGTCATTGGGAG-3’ and 5’-AGTTCTCCTCTGCACTCATAG-3’; *Ca_V_2.3* cDNA with 5’-GACTCTCATGTCACCACCGC-3’ and 5’-AGCCACTGGCATGTTCATCA-3’; and *beta-actin* with 5’-GCACCACACCTTCTACAATGAG-3’ and 5’-GAAGGTCTCAAACATGATCTGG-3’.

### Quantification and statistical analysis

Insulin secretion levels, ATP/ADP ratio of islets, glucose uptake of beta cells, actin remodeling, SFK activation, immunofluorescence intensities, protein expression profiles, chaperone binding capacities, proximity ligation assay, and blood sugar levels were subjected to Welch’s *t* test or one-way ANOVA. Ca^2+^ transients of beta cells, membrane depolarization of beta cells, Rho-family GTPase activation, and islet perifusion results were subjected to 2-way ANOVA. The movements of insulin granules were subjected to MSD analysis using the MSD Analyzer plugin (https://tinevez.github.io/msdanalyzer/) on the MATLAB platform (Tarantino *et al*., 2014). Statistical details of the experiments can be found in the figure legends, figures, results, and methods section. All error bars indicate the mean ± SEM.

## MOVIE LEGENDS

**Movie EV1. Exocytosis of a glucose-stimulated beta cells**

Time-lapse observations of insulin granule exocytosis in synapto.pHluorin-transduced primary beta cells of the indicated genotypes with a TIRF microscope. The whole movie corresponds to 1 min. Corresponding to Fig. 2A.

**Movie EV2. Mobility of insulin granules in primary beta cells.**

The mobility of insulin granules tagged with EGFP-phogrin in control primary beta cells, followed by that in cKO primary beta cells. The time lapse covers a period of 100 s approximately 30 min after glucose stimulation. Note that the mobility of insulin granules in cKO cells is decreased compared with that in control cells. Corresponding to Fig. 3C– H.

**Movie EV3. KIF5B-dependent dynamics of peripheral F-actin in beta cells.**

F-actin dynamics of control (CT) and cKO (KO) beta cells labeled by a *Lifeact-mCherry* transgene upon glucose stimulation. Scale bar, 5 μm. The duration of the movie corresponds to 28 min. Corresponding to Figs. 4A and EV2A.

**Movie EV4. KIF5B is largely dispensable for post-Golgi trafficking of the Ca_V_1.2 protein**

A BFA washout assay of Ca_V_1.2-EGFP in primary beta cells of the indicated genotypes. Bar, 5 μm. The Movie shows from 15 min to 119 min after the BFA washout. Corresponding to Fig. 6D.

**Movie EV5. Dynamics of cell bottom microdroplets containing Hsp90 and KIF5B**

Time-lapse recording in the bottom region of a primary wild-type mouse beta cell expressing tagRFP-Hsp90 (red) and KIF5B-EYFP (green). Scale bar, 1 μm. The duration of the movie corresponds to 464 s. Corresponding to Fig. 8A.

### Movie EV6. Dynamics of Hsp90 in an ER sheet

Multicolor time-lapse recording in the bottom region of a wild-type primary mouse beta cell, expressing tagRFP-Hsp90 (red) and mEmerald-Sec61β (green). Scale bar, 1 μm. The movie covers a period of 76 s. Corresponding to Fig. 8G.

---

**Figure EV1.**
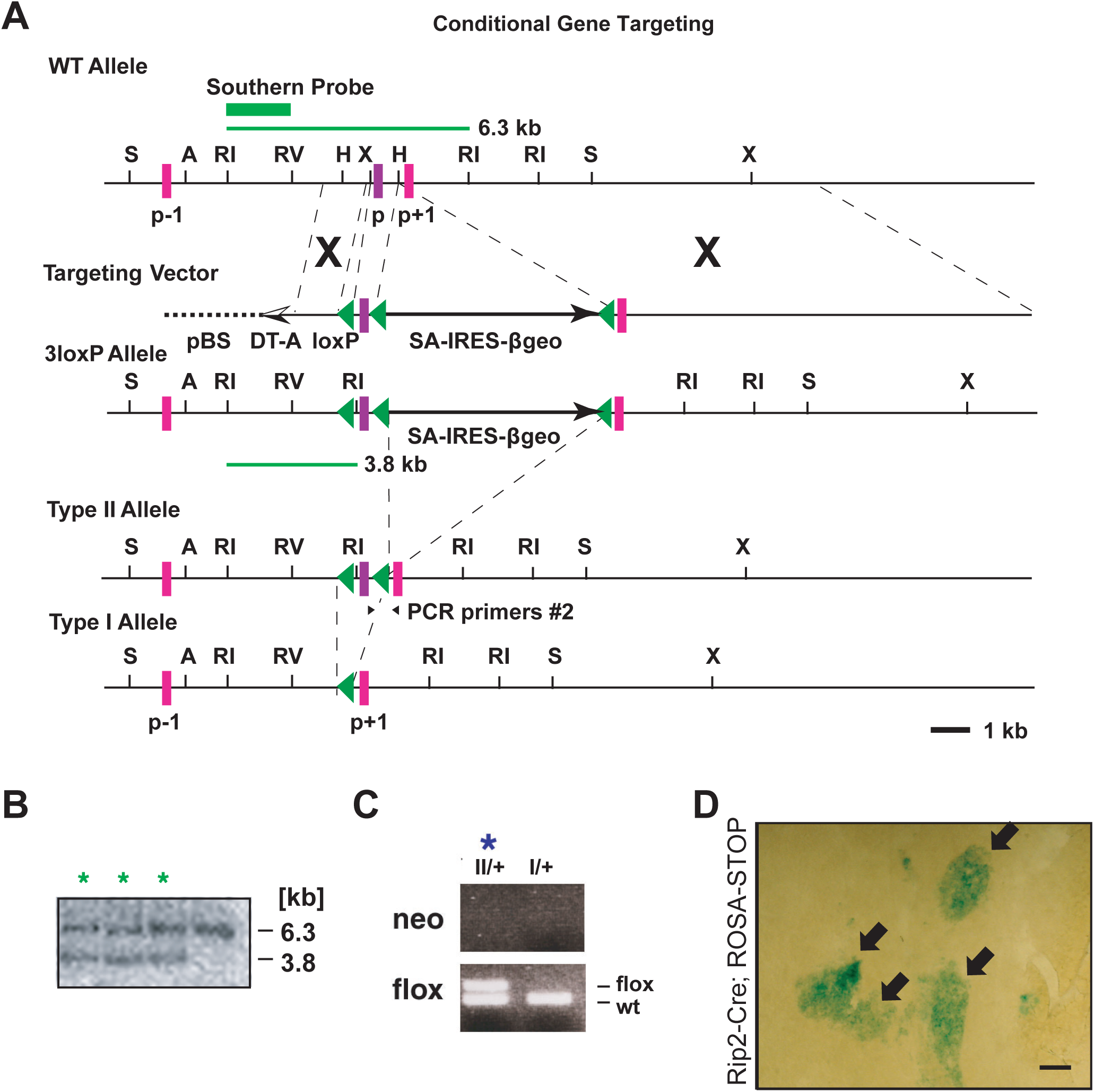
Conditional knockout of mouse *Kif5b* gene. (A–D) Establishment of beta-cell-specific *Kif5b* gene conditional knockout (cKO) mice, represented by a gene targeting strategy in mouse ES cells (A), Southern blotting screening for homologous recombinants (B; asterisks); genotyping PCR for the floxed allele (C: asterisk); and characterization of Rip2-Cre activity in a pancreas section detected by a LacZ reporter, ROSA-STOP mice (D). p, the 74 bp P-loop exon flanked by *loxP* sites (green triangles). S, *SalI*; A, *ApaI*; RI, *EcoRI*; RV, *EcoRV*; H, *HindIII*; X, *XbaI*. Arrows in D, specific Cre/*loxP* recombination sites in pancreatic islets of a *Rip2-Cre ROSA-STOP* double heterozygous mouse. Bar, 100 μm. Corresponding to Figure 1A–C.

**Figure EV2.**
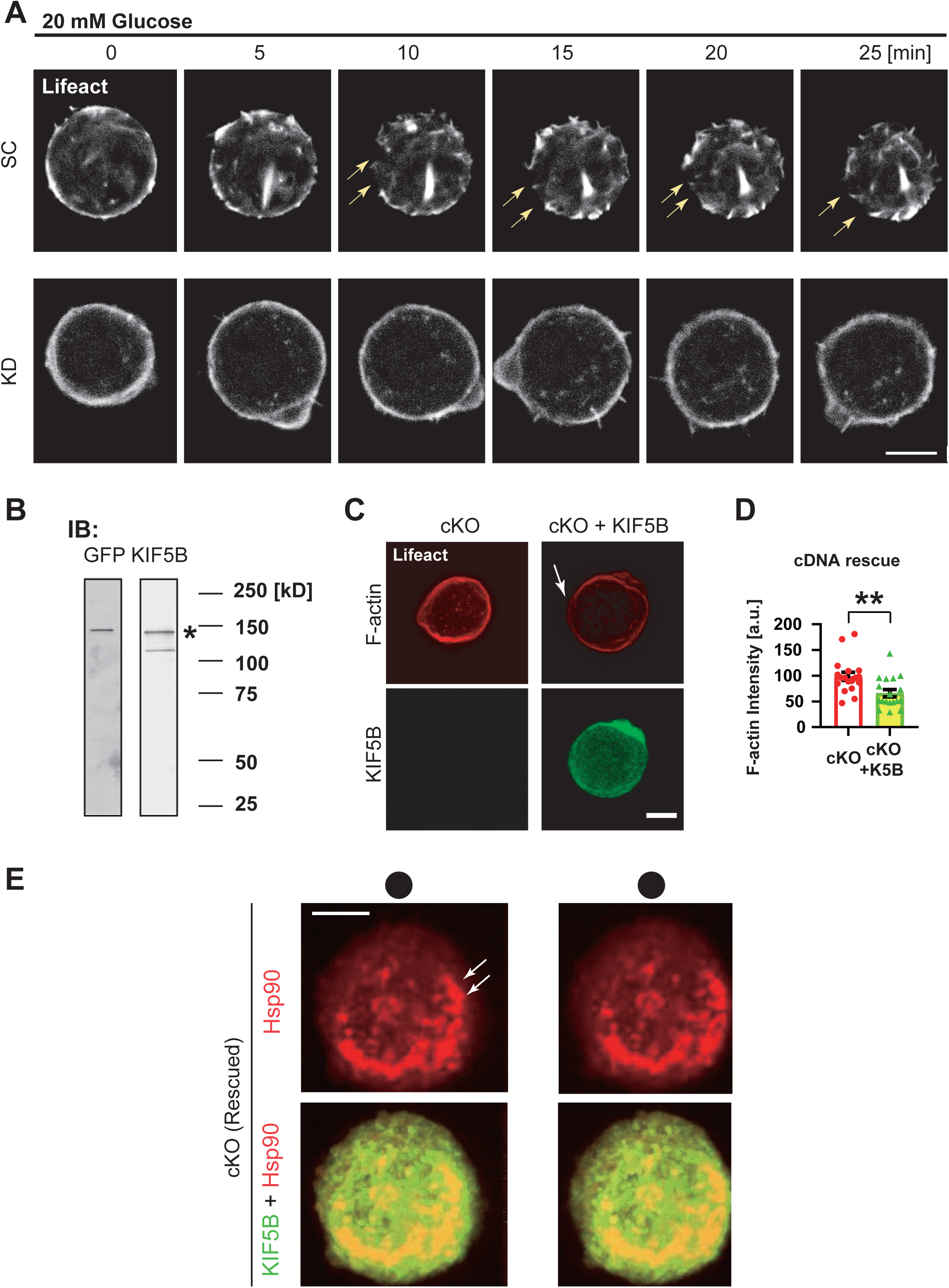
KIF5B facilitates cortical actin remodeling. (A) Time-lapse study of glucose-stimulated actin remodeling of primary beta cells from *Lifeact-mCherry* transgenic mouse pancreas, transduced with scrambled control (SC) or KIF5B-knockdown (KD) miRNA expression vectors. Scale bar, 5 μm. Corresponding to Fig. 4A and Movie EV3. (B–D) Rescue study of the glucose-stimulated actin remodeling in cKO primary beta cells by transducing KIF5B-EYFP, represented by immunoblotting of the expressed proteins in Ins1 cells using a mouse anti-GFP antibody and a rabbit anti-KIF5B antibody (B), *Lifeact-mCherry* transgene labelling (C), and F-actin quantification (D). Scale bar, 5 μm. Asterisks in B, bands for tagged KIF5B. Arrow in C, actin remodeling. \*\**p* < 0.01, Welch’s *t* test; n = 18–20. Corresponding to Fig. 4A. (E) Stereoscopic fluorescence microscopy of a cKO primary beta cell expressing tagRFP-Hsp90 (red) and KIF5B-EYFP (green). Scale bar, 5 μm. Corresponding to Fig. 8A.

